# Virus-host coexistence in phytoplankton through the genomic lens

**DOI:** 10.1101/513622

**Authors:** Yau Sheree, Marc Krasovec, Stephane Rombauts, Mathieu Groussin, L. Felipe Benites, Emmelien Vancaester, Jean-Marc Aury, Evelyne Derelle, Yves Desdevises, Marie-Line Escande, Nigel Grimsley, Julie Guy, Hervé Moreau, Sophie Sanchez-Brosseau, Yves van de Peer, Klaas Vandepoele, Sebastien Gourbiere, Gwenael Piganeau

## Abstract

Phytoplankton-virus interactions are major determinants of geochemical cycles in the oceans. Viruses are responsible for the redirection of carbon and nutrients away from larger organisms back towards microorganisms via the lysis of microalgae in a process coined the ‘viral shunt’. Virus-host interactions are generally expected to follow ‘boom and bust’ dynamics, whereby a numerically dominant strain is lysed and replaced by a virus resistant strain. Here, we isolated a microalga and its infective nucleo-cytoplasmic large DNA virus (NCLDV) concomitantly from the environment in the surface NW Mediterranean Sea, *Ostreococcus mediterraneus*, and show continuous growth in culture of both the microalga and the virus. Evolution experiments through single cell bottlenecks demonstrate that, in the absence of the virus, susceptible cells evolve from one ancestral resistant single cell, and vice–versa; that is that resistant cells evolve from one ancestral susceptible cell. This provides evidence that the observed sustained viral production is the consequence of a minority of virus-susceptible cells. The emergence of these cells is explained by low-level phase switching between virus-resistant and virus-susceptible phenotypes, akin to a bet hedging strategy. Whole genome sequencing and analysis of the ~14 Mb microalga and the ~200 kb virus points towards ancient speciation of the microalga within the *Ostreococcus* species complex and frequent gene exchanges between prasinoviruses infecting *Ostreococcus* species. Re-sequencing of one susceptible strain demonstrated that the phase switch involved a large 60 Kb deletion of one chromosome. This chromosome is an outlier chromosome compared to the streamlined, gene dense, GC-rich standard chromosomes, as it contains many repeats and few orthologous genes. While this chromosome has been described in three different genera, its size increments have been previously associated to antiviral immunity and resistance in another species from the same genus. Mathematical modelling of this mechanism predicts microalga–virus population dynamics consistent with the observation of continuous growth of both virus and microalga. Altogether, our results suggest a previously overlooked strategy in phytoplankton–virus interactions.

## Introduction

NCLDVs infect phytoplankton lineages across the entire eukaryotic tree of life: Chlorophyceae, Cryptophyceae, Dinophyceae, Mamiellophyceae, Pelagophyceae, Prymnesiophyceae, and Raphidophyceae [1][2][3]. While cultivation-free metagenomic approaches have enabled the estimation of the astounding abundance and genetic diversity of NCLDVs in planktonic communities [4], there are only a handful of cultivable NCLDV– microalgae systems available to study the mechanisms underlying their interactions [5]. Indeed, precise estimations of virulence and resistance parameters, and how they co-evolve, are required for modelling the flow of energy and matter in the oceans [6]. Resolving the mechanisms underlying these interactions and their significance to bloom development represents one of the major challenges for future research on ocean ecosystems dynamics under climate change [7][8]. Environmental surveys estimated NCLDV to be present at 10^4^.ml^−1^ in the photosynthetic zone [4], for which the majority were affiliated to prasinoviruses, a monophyletic group within the Phycodnaviridae infecting Mamiellophyceae, a class of globally distributed marine picophytoplankton [9]. Within the Mamiellophyceae, the genus *Ostreococcus* and its prasinoviruses represent ideal, ecologically relevant, model systems for focused study of host–virus interaction mechanisms and dynamics [10]. Notably, genome data is available for 16 *Ostreococcus* strains, 12 prasinoviruses [11][12][10], and fostered the discovery of one structurally unstable[13] and hypervariable [14] small outlier chromosome implicated in the resistance of the microalga to prasinoviruses. *Ostreococcus mediterraneus* is the most recently described *Ostreococcus* species and gene phylogeny of the 18S rRNA indicated it is the earliest branching member of the genus [15]. Preliminary analysis of the complete genome sequence of *O. mediterraneus* serendipitously revealed the genome of a novel prasinovirus, hereafter named OmV2 for *Ostreococcus mediterraneus* Virus 2. As we could observe the presence of OmV2 particles in the present cultures and trace back its presence in the culture for a decade, we decided to take advantage of this novel microalga–NCLDV system to unravel the mechanisms of microalga-virus coexistence. Unlike bacteriophages, which may integrate into their bacterial hosts as prophages, and develop complex lysogeny-lysis switch strategies [16], all known viruses thus far infecting unicellular algae replicate by cell lysis [17]. However, acquisition of viral resistance after initial lysis by viruses has been reported in many phylogenetically diverse microalgal–NCLDV systems [18], but the origin of continuous production of infective viruses in dividing microalga cultures has remained unsolved. Importantly, “coexistence” of microalgae and viruses is here defined as virus production by healthy microalga cultures isolated from one single cell, and thus isogenic cultures, except for the presence of spontaneous mutations. Virus and microalga coexistence has been first reported in the freshwater green alga *Chlorella variabilis* N64A infected by PBCV-1, and this phenotype was coined a “carrier” state [19]. Coexistence was observed later in the haptophyte *Emiliania huxleyi* upon infection by EhV-99B1 over 12 days [18], and subsequently in *O. tauri* upon infection by OtV5 over 18 days [20]. More recently, a study of the population dynamics of the freshwater green alga *Chlorella* and PBCV-1 [21] reported stable coexistence evolved after 30 days post-infection, which was maintained until the end of the experiment 45 days later.

Here, we analyze the genomes of *O. mediterraneus* and OmV2 as a microalga–virus system that was isolated as a coexisting system, rather than upon the selection of resistant cells after an initial lytic event. We investigate the origin of the viral production and test for resistance–susceptible phenotype switching of the microalga by experimental evolution. Mathematical modelling of this mechanism enabled predictions of the effect of such a switch on population dynamics of this system, and comparison with the dynamics observed in experimental conditions.

## Materials and Methods

### Cultures and growth conditions

The isolation of *O. mediterraneus* strains from the environment was described previously [15]. All strains were maintained in liquid L1 medium (NCMA, Bigelow Laboratory for Ocean SEvociences, USA) without addition of Na_2_SiO_3_, prepared using autoclaved seawater from offshore Banyuls Bay (MOLA station: 42°27’11"N, 3°32’36"E) diluted 10% with MilliQ water and filtered prior to use through 0.22 μm filters, under a 12:12 hour light:dark regime. Incubation temperature was 15°C for measuring growth curves in population dynamics experiments and 20°C for all other experiments. The strains used in this study, the genomic and transcriptomic data obtained for them are listed in Supplementary Table 2.

### PFGE and in-gel hybridisation

PFGE and in-gel hybridization enabled estimation of chromosome sizes in different strains and testing for the presence of the virus genome. PFGE and in-gel hybridisation was performed as previously described [22][23] with the following modifications: 8.7 × 10^7^ cells in exponential growth phase were harvested by centrifugation (8000 *g* for 20 min) and resuspended in 100 μl of TE buffer (10 mM Tris-HCl, 125 mM EDTA, pH 8), then embedded in plugs by mixing with an equal volume of molten low melting point agarose (1% in TE buffer precooled to 45°C). PFGE was performed using the CHEF-DR III (Bio-Rad) system at 3 V cm^−1^ at 14°C with 120° pulse angle for 45 h with a switch time of 90 s and followed by 27 h at a switch time of 140 s. PCR (GoTaq, Promega) was used to amplify specific regions of the SOC, BOC, OmV2 and the 18S rRNA gene (see Supplementary Table 3 for primer sequences) for use as DNA probes with the following conditions: initial denaturation for 2 min at 94°C; 30 cycles of denaturation for 30 s at 94°C, annealing for 30 s at 55°C, and extension for 45 s at 72°C; final extension for 5 min at 72°C.

### Transmission electron microscopy

RCC2590 culture in exponential growth phase was prepared for transmission electron microscopy (TEM) as previously described (Derelle et al. 2008). Images were captured of individual cells and cells deemed to be visibly infected if clear virus-like particles were clearly present within the cell.

### Genome sequencing and assembly

Genomic DNA was extracted from *O. mediterraneus* RCC2590 as described by [24]. Genome library preparation, sequencing, assembly and optical mapping were performed by GENOSCOPE (http://www.genoscope.cns.fr/spip/). Two libraries were prepared and sequenced using the Illumina Hi-Seq platform; a paired-end (PE) read library of insert size 272 bp and a mate-pair (MP) library of 5 230 bp insert size. DNA sequence reads were assembled using ALL-PATHS-LG assembler [25]. At this stage, 99 scaffolds were obtained that aligned with eukaryotic sequences. Optical mapping was performed to further place the *O. mediterraneus* scaffolds in their correct order on the chromosome. This resulted in 19 optically mapped scaffolds corresponding to nuclear chromosomes and several low GC scaffolds corresponding to fragments of the SOC, organelles and OmV2.

The organelles were assembled using ABySS from the Illumina Hi-Seq PE reads using the MP library for scaffolding (abyss-pe, lib = ‘PE’, mp = ‘MP’, k = 85, n = 10). Organelle genome completeness was evaluated by alignment to the complete organelle sequences of *O. tauri* [26]. The two large contigs of the OmV2 genome obtained from the Illumina sequencing were scaffolded together manually using the assembly and visualisation tools available from the Geneious software [27].

To resolve the highly repetitive SOC sequence, genomic DNA from RCC2590 and the evolved line MA3 were re-sequenced with the SMRT Technology PacBio RS II by the GetPlaGe sequencing platform in Toulouse, France as previously described [14]. The raw data was assembled using HGAP3 [28]. The PacBio assemblies of were checked manually with assembly and visualisation tools available from the Geneious software to identify the scaffolds corresponding to the SOC. Both RCC2590 and MA3 SOCs were assembled from telomere to telomere, the former comprising 5 scaffolds and the latter a single scaffold.

### Genome annotation

The RCC2590 genome was annotated using the annotation strategy described in [29]. Briefly, gene models were predicted using Eugene (release 4.1; http://eugene.toulouse.inra.fr/) [30]. Transcript information was provided to the gene-caller as follows: one paired-end Illumina RNAseq dataset was assembled into transcript contigs using a DBG assembly approach with high Kmer sizes to boost specificity during the assembly. Six *O. mediterraneus* strains (Supplementary Table 2) had their transcriptome sequenced as part of the Marine Microbial Eukaryotic Transcriptome Sequencing Project (MMETSP) [31]. The resulting transcriptomes are referenced in Supplementary Table 2 with their respective MMETSP identifiers.

The obtained set of contigs was cleaned from any contigs smaller that 300 bp, completed with EST sequences collected from NCBI (>300 bp), and mapped on the genomic scaffolds by combining BLASTN (for rapid detection on regions of interest) and GenomeThreader [61] (for fine mapping, taking splice sites into account) with parameters set to map at least 98% of the transcripts sequence on the regions as reported by blastn. Besides, RNAseq was also mapped directly onto the genome using HISAT2 [33] and junctions were extracted from the BAM alignments with regtools. The obtained junction instances where counted and grouped based on the intron coordinates that was spanned by a read. This count (or coverage) was used to filter for junctions confirmed by minimum 3 reads. Functional descriptions were inferred from filtered best-BLAST-hits and combined with protein domains obtained with Interpro. This automatic annotation was curated using the ORCAE portal [34]. All predicted genes were manually inspected and from the 8483 protein coding genes, 1142 have been modified and 59 discarded, resulting in a final set of 8424 protein coding genes. The functional description was manually edited for 662 genes.

Gene prediction in OmV2 was performed using Glimmer implemented within Geneious [35]. Functional description of predicted OmV2 genes was done using BLASTX against the NR database (from December of 2017) with default parameters and against the Pfam version 31.0 database (http://pfam.xfam.org/). tRNAs were predicted using the tRNAscan-SE 2.0 web server [36].

### Comparative genome analysis

The genome and predicted genes from *O. mediterraneus* RCC2590 were loaded into a custom instance of pico-PLAZA [37] containing 38 eukaryotic genomes (including Metazoa, Fungi, Chlorophyta, Embryophyta, Rhodophyta, Haptobionta and Stramenopiles, Supplementary Table 5). Following an ‘all-versus-all’ BLASTP [38] (version 2.2.27+, e-value threshold 10^−5^, max hits 500) protein sequence similarity search, both TribeMCL [39] (version 10-201) and OrthoMCL [40] (version 2.0, mcl inflation factor 3.0) were used to delineate gene families and subfamilies, respectively.

Phylogenomic analysis of the *O. mediterraneus* nuclear genome was performed by concatenating the amino acid sequence from 188 single copy orthologs by TribeMCL within the Chlorophyta genomes (*O. mediterraneus, O. tauri, O. lucimarinus, O. sp RCC809, Micromonas pusilla strain CCMP1545, Micromonas sp RCC299, Bathycoccus prasinos, Arabidopsis thaliana, Chlamydomonas reinhardtii, Coccomyxa subellipsoidea C-169, Chlorella variabilis NC64A, Oryza sativa, Physcomitrella patens* and *Volvox carteri*) aligned using MUSCLE (version 3.8.31) [41]. This resulted in alignment of 148,028 positions of which 88,784 were informative. Similarly the phylogenetic topology across Mamiellales was determined by concatenating the alignment of 2,161 single copy genes, generated using MUSCLE, resulting in an alignment of 1,267,142 positions of which 444,911 were informative. The phylogenetic trees were generated using RaxML [42] (version 8.2.8) (model PROTGAMMAWAG) with 100 bootstrap replicates.

The phylogenetic profile of TribeMCL gene families (excluding orphans) and the inferred Mamiellales tree topology were provided to reconstruct the most parsimonious gain and loss scenario for every gene family using the Dollop program from the PHYLIP package (version 3.69) [43].

### Prasinovirus phylogenomics

85 putative orthologous prasinovirus genes were identified for phylogenetic analyses by tBLASTx of predicted OmV2 genes against a database of all genes from 15 fully sequenced prasinovirus genomes (e-value < 10^−4^, amino acid identity > 75% and alignment length >50) accepting as ‘core’ those orthologs present in all genomes. Each ortholog family was aligned using MAFFT [44] and phylogenetic trees were reconstructed using Bayesian inference with MrBayes [45] using a mixed model for AA with 4 chains of 10^6^ generations, trees sampled every 100 generations, and burnin value set to 20% of the sampled trees. We checked that standard deviation of the split frequencies fell below 0.01 to ensure convergence in tree search.

### Analysis of the evolution of GC content

To reconstruct genomic GC contents along the tree of *Ostreococcus*, we used an ancestral sequence reconstruction approach to reconstruct genes sequences at the codon level and compute their GC contents at the tree nodes. We included 3 outgroups (*B. prasinos*, *M. pusilla* CCMP1545 and *M. commoda* RCC299) in the reconstruction of ancestral GC contents of the four *Ostreococcus* lineages and selected all 2,161 single-copy genes universally present in these seven genomes as determined from the comparative genomic analysis.

We aligned all 2,161 universal single-copy protein families at the codon level with MACSE [46] using default parameters and trimmed alignments for badly aligned positions with BMGE [47] using a BLOSUM62 substitution matrix and other parameters set to their defaults values.

To both reduce computational burden and produce confidence intervals in estimates of ancestral GC contents, we employed a jackknife approach to subsample the 2,161 alignments and to build 100 concatenates containing each 100 randomly sampled genes. Each concatenate has an average size of 104,290 nucleotide positions (± 8,644), representing on average 34,763 codons, and so is expected to contain a sufficient amount of phylogenetic information to reconstruct ancestral GC contents. We further verified that these concatenates are faithful representations of each genome in terms of GC contents. We computed correlation coefficients between the average GC contents computed across all 2,161 genes and the GC contents of each of the 100 concatenates. We observed that all correlation coefficients were higher than 0.972, showing that these 100 jackknife sub-samples have the same properties than the original population of 2,161 genes, which allows us to infer ancestral genomic GC contents.

We first reconstructed the phylogenetic tree of each of the 100 concatenates to verify that each concatenate contains the same phylogenetic information on the relationships between the seven considered species. We built trees with FastTree (v. 2.1.8) [48] using a GTR model and the CAT approximation with 20 categories to model the variation of evolutionary rates across sites. Each phylogenetic reconstruction yielded a tree consistent with the reference species tree of the seven species. We used this topology to reconstruct ancestral sequences on all concatenates. We used the Yang and Nielsen (YN) substitution model for codons [49] with different base frequencies at the three codon positions (F3 × 4) to model codon evolution. As the seven species have genomic GC contents that vary from ~0.5 to ~0.65, the G+C composition has not evolved homogeneously in time, as was previously observed in many prokaryotic and eukaryotic lineages [50][51][52]. For this reason, we considered a non-homogeneous and non-stationary approach in which stationary frequencies are allowed to vary from one lineage to another on the species tree. Each branch of the species tree is assigned its own set of parameters of the YN model and the stationary codon frequencies at the root of the species are also free to vary. We used Maximum Likelihood (ML) and the bppML program [53][54] to estimate the parameters of this non-homogeneous model and the branch lengths of the tree. Using the ML values for all these parameters, we reconstructed ancestral sequences with the bppAncestor program [53], using the marginal ASR approach [55]. At a given internal node of the species tree, the codon state having the maximum posterior probability was inferred as the putative ancestral state. We then computed the ancestral global GC content as well as the ancestral GC content of all codon positions from these reconstructed ML ancestral sequences. We obtained a distribution of ancestral GCs for each ancestor from which we computed the mean and the 95% confidence interval.

### Population dynamics of virus and microalga

Microalgae and viruses were counted over 13 days by flow cytometry and plaque assay from triplicate 20 ml cultures in tissue culture flasks at 15°C in a 12 h light:12 h dark regime. Cultures of RCC2590 and MA3 were inoculated to have a starting density of 10^4^ cells.ml^−1^. Additionally, we passed the RCC2590 culture through a 0.2 μm filter to remove cells retaining free virions in the medium and kept this filtrate in the same conditions as the cultures to estimate viral particle stability over the duration of the experiment. For flow cytometry counts of cells and viruses, 250 μl was sampled every two days, fixed for 15 min in the dark with a final concentration of electron microscopy grade glutaraldehyde of 0.25% and Pluronic F68 of 0.01%, flash frozen in liquid nitrogen and stored at −80°C until analysis as prescribed in [56]. Cell and viral particle counts were performed with the BD FACSCanto II (Becton Dickinson) where the flow rate over 1 min at a low rate was determined using triplicate measurements of BD Trucount Tube beads. Cells were counted from fixed samples using red autofluorescence and side scatter. Viral counts were performed on fixed samples after rapid thawing at 40°C, diluting as necessary in filtered seawater (< 0.02 μm), staining for 10 min at 80°C with SYBR Green I (Molecular Probes) (0.5× final concentration of commercial stock) and setting the trigger to measure green fluorescence and side scatter. To measure the concentration of infective OmV2 in the RCC2590 culture medium by plaque assay, 1 ml was sampled from the cultures every two days, centrifuged at 25,000 *g* for 10 min at 15°C to remove cells and the supernatant serially diluted with L1 medium. Plaque assay was performed with the diluted supernatant as described in [39] using the OmV2-susceptible *O. mediterraneus* strain MA3 as the host.

### Experimental assessment of the Resistant to Susceptible phase switch

Virus-free *O. mediterraneus* lines were obtained by end point dilution and virus absence was confirmed by PCR using degenerate primers for the viral DNA polymerase and Major Capsid Protein gene (Supplementary Table 3). We started 204 clonal cultures from one single cell in 1 ml microwells as described in [57] and tested for virus susceptibility after 7 days. Virus susceptibility was tested using cell-free virus producing culture (VPC) medium.

To obtain cell-free VPC medium (i.e. separate potential virions from cell lysates), medium from growing or lysed cultures was centrifuged at 25,000 *g* for 10 mins at room temperature and the supernatant filtered (<0.45 μm). To test if a cell line was susceptible to lysis, VPC medium was introduced into the growing test culture, with a control culture grown in parallel inoculated with the same volume of sterile medium, and visually inspected for lysis up to five days after inoculation. This visual inspection detected when most cells are lysed and thus the ancestral cell was inferred to be susceptible. The validity of this inference relied on two assumptions: a low switch rate between the susceptible and resistant phenotype, and a low difference in growth rates between the resistant and susceptible phenotypes.

### Mathematical modelling of the host-virus population dynamics

The model considered a population made of susceptible and resistant microalga (*S*_*t*_, *R*_*t*_) and their viruses (*V*_*t*_) at time *t*. The changes in microalgae and viruses numbers from one day (t) to another (t+1) were modelled according to a simple representation of the algae and virus life-cycles and interactions with following assumptions: i) susceptible and resistant microalgae produce new microalgae at rate *a*_*s*_ and *a*_*r*_, ii) microalgae born from a susceptible (resistant) to switch to a resistant (susceptible) phenotype at a fixed rate *e*_*R*_(*e*_*S*_), iii) encounters between susceptible algae and virus to occur randomly at a rate *c*, iv) each infected microalga produced *v* viruses before death, and v) viral particles survived at a rate *s*_*V*_. The population dynamic of *R*, *S* and *V* can then be described by following equations:

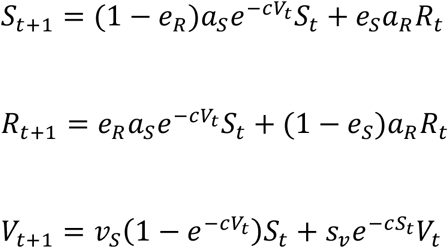

where 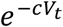 is the probability drawn in Poisson distributions that a susceptible algae encounters no virus or that a viral particle encounters no susceptible algae to infect. This system was investigated analytically and numerically using the R software.

## Results

### Genomic features of *O. mediterraneus* and OmV2

The *O. mediterraneus* RCC2590 genome assembly comprised 20 large scaffolds, in agreement with the number and sizes of chromosomes estimated from Pulsed Field Gel Electrophoresis (PFGE) (Figure 1). Probes designed to unique regions of three chromosomes: the big outlier chromosome (BOC), carrying the candidate sexual mating type locus [58][14], the predicted small outlier chromosome (SOC), involved in viral resistance [13][14], and chromosome 17, carrying the 18S rRNA gene, hybridised to chromosomal bands in the gel corresponding to their assembled sizes. Sixteen scaffolds were bordered by two telomeric sequences, providing further validation that this genome assembly comprised a majority of complete chromosome sequences. Complete circular assemblies were also obtained for the chloroplast and the mitochondrion, which were similar to other *Ostreococcus* species in size, topology, gene content and the presence of segmental duplications [59]. *O. mediterraneus* also displayed a similar number of chromosomes and genes, as well as the hallmarks of genome streamlining seen in other Mamiellophyceae genomes, including relatively small overall genome size and high coding density. Two striking features of this genome were the presence of the largest SOC so far recorded, which was more than three times the average size of other sequenced Mamiellophyceae, and a relatively low GC content (56%) compared to other *Ostreococcus* species (Table 1). Based on an alignment of protein coding regions between *O. tauri* and *O. mediterraneus*, all chromosomes, except for the SOC and the putative mating locus of the BOC, shared a high degree of co-linearity (Supplementary Figure 1).

**Figure 1.**
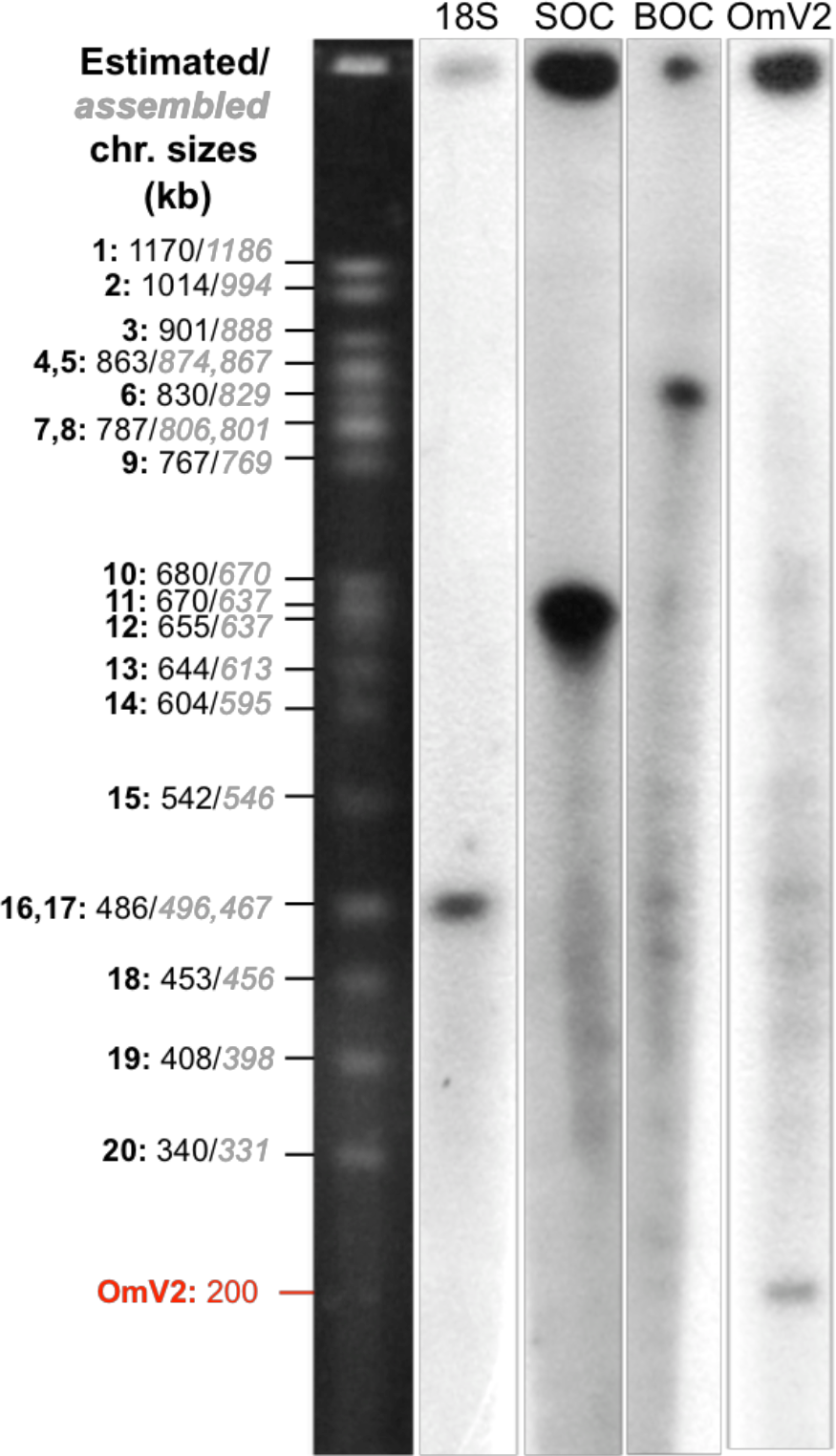
PFGE of the chromosomes of *O. mediterraneus* RCC2590. The estimated chromosome number and sizes are inferred from band intensity and mobility, respectively, and chromosome sizes from the assembled genome are indicated left black background. Hybridizations to the PFGE with radioactive probes specific to the 18S rRNA gene, SOC, BOC and OmV2 regions are indicated on the right in a white background. Estimated prasinovirus OmV2 size is shown in bold red. The topmost band present in all lanes is non-specific hybridization to unmigrated DNA remaining in the loading wells. Probes were generated by PCR using primer pairs listed in Supplementary Table 3.

**Table 1.** Comparison of the genomic features of sequenced Mamiellophyceae.

|  | *O. mediterraneus* | *O. tauri* | *O. lucimarinus* | *Ostreococcus sp. RCC809* | *B. prasinos* | *M. pusilla*<br>CCMP1545 | *M. commoda*<br>RCC299 |
| --- | --- | --- | --- | --- | --- | --- | --- |
| **Size (Mb)** | 13.86 | 12.91 | 13.20 | 13.30 | 14.95 | 21.9 | 20.9 |
| **No. chromosome** | 20 | 20 | 21 | 20 | 19 | 19 | 20 |
| **GC content (%)** | 56 | 60 | 60 | 60 | 48 | 65 | 64 |
| **Genes** | 7.85 | 7.749 | 7.603 | 7.492 | 7.847 | 10.587 | 10.286 |
| **genes/kb** | 0.60 | 0.62 | 0.58 | 0.56 | 0.52 | 0.48 | 0.49 |
| **Mono-exonic genes (%)** | 77 | 81 | 79 | 81 | 86 | 47 | 63 |
| **BOC size (kb) [chr no.]** | 874 [4] | 1,217 [2] | 895 [2] | 822 [2] | 663 [14] | 2172 [61] | 2053 [61] |
| **Candidate mating type** | *plus* | *minus* | *minus* | *plus* | - | - | - |
| **SOC size (kb) [chr. no.]** | 637 [12] | 312 [14] | 150 [18] | 102 [18] | 146 [62] | 242 [61] | 195 [61] |
| **Mitochondrion size (bp)** | 49.312 | 44.237 | - | 43.168 | 43.614 | - | 47.425 |
| **Chloroplast size (bp)** | 74.215 | 71.666 | - | 67.238 | 72.700 | - | 72.585 |

A linear 192 kb genome of OmV2 with mean GC content 40.7% was assembled, encoding 221 predicted genes, similar to all other sequenced prasinoviruses [11][60][61][12]. Visualisation of the DNA extraction of *O. mediterraneus* cultures by PFGE and subsequent hybridisation with OmV2 specific probes showed a band at ~200 kb (Figure 1), consistent with the assembled genome size and a linear structure. Importantly, this hybridisation demonstrated that the viral genome was not integrated into one of the *O. mediterraneus* chromosomes. The abundance of OmV2 in the culture was so low as not to be visually detectable as a band by ethidium bromide staining (indicating <1 ng of OmV2 in the PFGE). The coverage of the OmV2 genome was ~0.5× that of the microalga nuclear genome, suggesting that, at the time of sequencing, OmV2 genome copies were relatively more abundant.

### Evolution of genome architecture in *Ostreococcus*

The phylogenetic position of *O. mediterraneus* at the earliest diverging branch within the *Ostreococcus* genus was confirmed by phylogenomic analysis of 188 single-copy gene families shared among all Chlorophyta (Supplementary Figure 2) and from 2,161 single copy Mamiellales orthologues (Figure 2). Comparative protein family analysis was conducted on the fully sequenced Mamiellales genomes to determine if protein families have been lost, gained or specifically expanded in *O. mediterraneus*. This showed all *Ostreococcus* species have lost 261 of the 5096 protein families present in all Mamiellales (Figure 2), which represent probable ancestral Mamiellales functions and thus reflect the general trend of genome reduction in *Ostreococcus* (Table 1). Moreover, *O. mediterraneus* showed few significantly expanded or reduced protein families relative to the other *Ostreococcus* species. Nonetheless, *O. mediterraneus* has retained relatively more of these ancestral protein families, of which 72 are uniquely present in *O. mediterraneus* and lost in the other *Ostreococcus* species. This is further support of *O. mediterraneus* being the earliest branching *Ostreococcus* species.

*O. mediterraneus* has a low genomic GC content compared to other *Ostreococcus* species, raising the question of whether it descended from a high GC-content ancestor and experienced a shift towards a lower GC composition, or whether it descended from a low GC-content ancestor and that an increase in GC occurred in the rest of the *Ostreococcus* clade. To discriminate between these two alternative hypotheses, we inferred the ancestral genomic GC content for all ancestors of the species tree using ancestral sequence reconstruction techniques. Because current GC-contents vary between 0.5 and 0.65, we reconstructed ancestral sequences with a non-homogeneous codon substitution model to allow each lineage to vary in global nucleotide compositions. We observed that the last common ancestor (LCA) of the *Ostreococcus* clade had a GC-content of ~0.57, similar to that of *O. mediterraneus*, and lower than that of *O. tauri*, *O. lucimarinus* and *Ostreococcus* sp. RCC809 (~0.60) (Figure 2). This global trend is consistent across all codon positions, although to a much lesser degree for the highly constrained second codon position. The trend observed at the genomic level is mostly driven by the third codon position, where the differences between the GC-content of the *Ostreococcus* LCA and the GC-content of the other *Ostreococcus* species is exacerbated (sharp increase from 0.66 to 0.78) (Supplementary Figure 3). Interestingly, GC content evolution with Mamiellophyceae reflects two independent episodes of GC increase, one within *Ostreococcus*, and the other in the branch leading to the *Micromonas* lineage (Figure 2).

**Figure 2.**
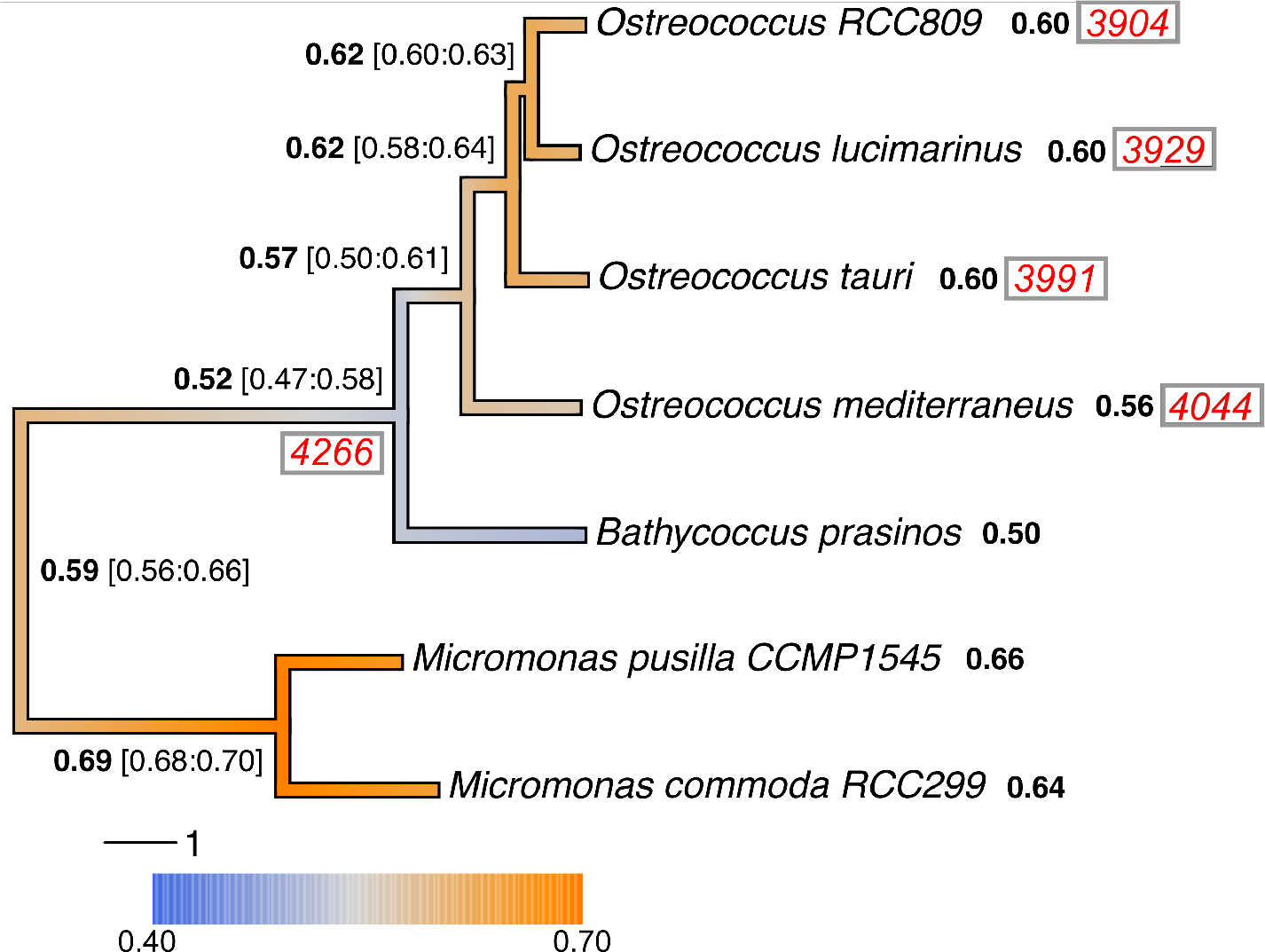
Phylogeny, reconstructed ancestral global GC contents and protein families of fully sequenced Mamiellales genomes. The phylogenetic tree was calculated from random subsampling of 2,161 Mamiellales single copy orthologous gene alignments. The branch colours correspond to the estimated changes in GC over time. The average genome wide GC contents of extant genomes are shown after the species in bold and the ancestral estimates shown at the nodes with ranges in square brackets (see *Methods* for the GC evolution estimates). The total number of protein families common to *Bathycoccus* and *Micromonas* are shown in red italics at the node leading to those genera and the number of those orthologs present in extant *Ostreococcus* are shown after the species names.

### OmV2 gene genealogies tell intertwined microalga-prasinovirus evolutionary histories

Whole genome alignment of OmV2 against the model *Ostreococcus* virus OtV5 [11], revealed large syntenic regions and overall conservation of gene order, interspersed with virus specific gene gains and losses (Figure 3A, Supplementary Table 1). Phylogenetic analysis of the widely used NCLDV taxonomic marker, the DNA polymerase B (PolB) gene, clearly placed OmV2 with viruses infecting *Ostreococcus* spp. within the *Prasinovirus* (Figure 3B). It also revealed that the PolB topology clustered viruses infecting the microalgae from the same genus together, and although viruses of the same clade are generally specific to the same host species, some viral clades contain members that were isolated from different species. This suggests that closely-related viruses are constrained to genetically similar hosts, but switching to related host species has probably occurred in prasinovirus evolutionary history [62]. To explore the evolutionary relationship of OmV2 with other prasinoviruses, we identified 85 core orthologous proteins shared between the 15 complete prasinovirus genomes. Phylogenies computed for each of these sequences showed that most orthologs supported the monophyly of *Ostreococus* viruses, with only a few clustering with MpV1 (Figure 3C). However, the tree topologies within the *Ostreococcus* viruses were not all congruent. The majority (36%) of OmV2 core genes were in the same clade as OmV1, OtV1 and OtV5 (Figure 3B) as compared to 8% of prasinovirus orthologs supporting its more basal position. This variability in gene evolutionary histories could be the consequence of recombination between related viral clades [63], as well as independent horizontal gene transfers from different microalga hosts into the virus genome [64].

**Figure 3.**
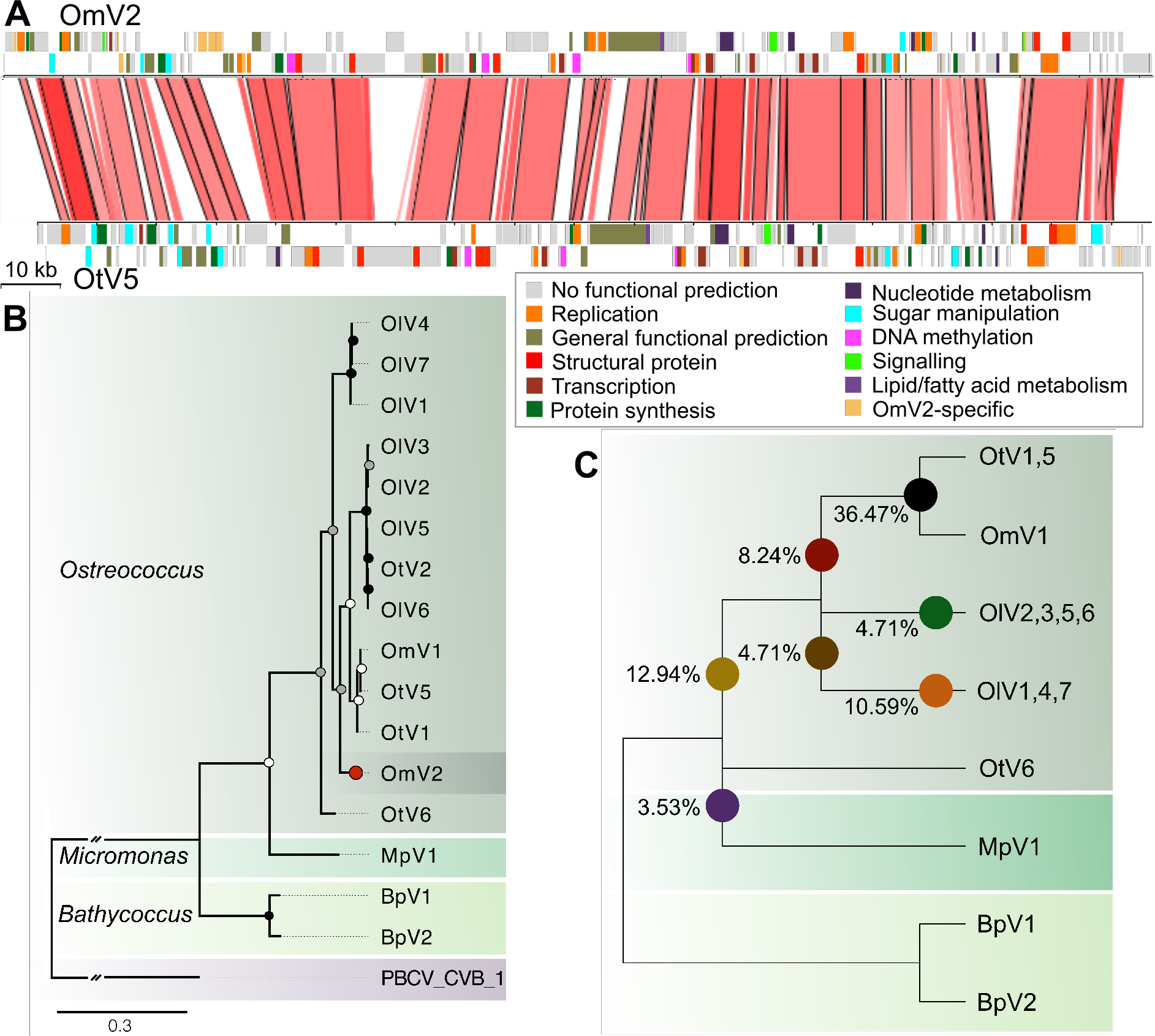
Phylogenomic analysis of prasinovirus OmV2. **(A)** Whole genome alignment of OmV2 compared to the model *Ostreococcus* virus, OtV5. Genomic regions with 60-100% nucleotide identity are joined by red blocks with higher identity indicated by darker red. Genes are coloured according to functional groups. **(B)** Maximum likelihood phylogenetic tree of amino acid sequences of the DNA polymerase B gene from prasinoviruses infecting the genera *Ostreococcus*, *Micromonas* and *Bathycoccus*. The tree was rooted with *Paramecium bursaria chlorovirus* (PBCV1) (Chlorovirus) with the connecting branch truncated for display. The red circle highlights the position of OmV2. Circles at the nodes represent bootstrap support of 50-70% (white), 70-90% (grey) and 90-100% (black). The scale bar shows substitutions per site. **(C)** Cladogram representing the relationships between the fully sequenced prasinovirus clades from 85 core genes. Coloured circles mark the branching positions of the percentage of the OmV2 orthologs.

### Production of virions is the consequence of lysis of a subset of microalgae

Population dynamics of the microalga and the virus was followed by flow cytometry during two weeks. Virus concentration was estimated by two methods. First by flow cytometry and second by plaque assays, taking the number of lysis plaques as a proxy of the number of infective particles. The latter method aims at estimating the number of infectious viruses, while the former estimates the number of total viral particles in the culture medium. The logarithm of microalga concentration, *C*, increases linearly with time, *t*, in RCC2590 culture (Figure 4, ln(*C*)= 0.557×*t*+7.559, *R*^2^=0.94). Despite variation in the estimation of the number of viruses between replicates and between the two methods, the number of viruses significantly increases with the number of microalga, providing evidence of a continuous production of viruses in the culture.

**Figure 4.**
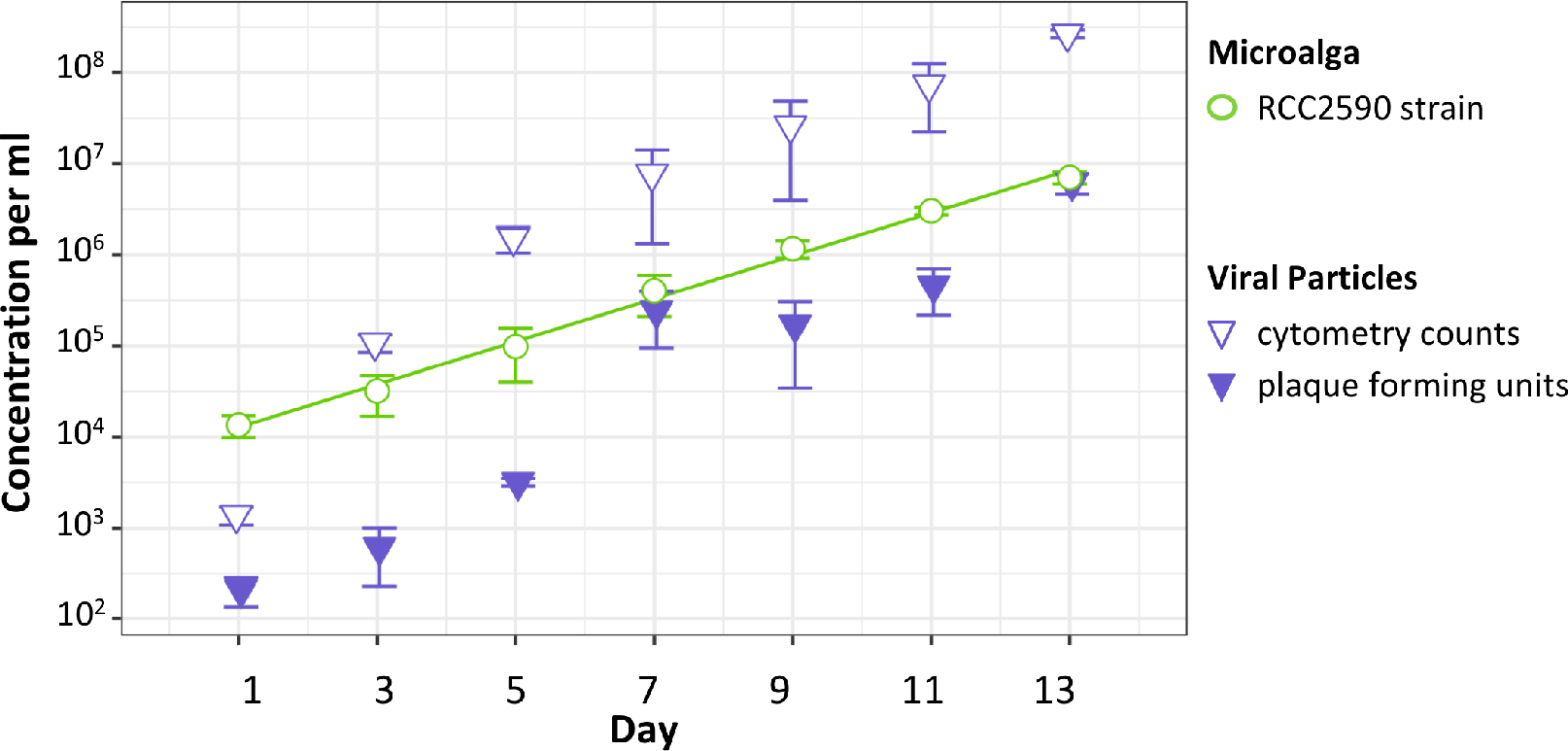
Population dynamics of microalga (circles) and prasinovirus (triangles) concentrations over 14 days in the *O. mediterraneus* RCC2590 culture. The line is fitted via ordinary least square regression.

To gain insight into how OmV2 was being produced, we performed visual inspection of transmission electron microscopy (TEM) thin sections of RCC2590 cultures for signs of viral infection. Indeed, the majority of cells had typical morphology (Figure 5A) while 0.9% of cells (5 of 573 individuals) contained icosahedral particles of approximately 110 nm in diameter within the cytoplasm (Figure 5B and C). Viral particles could also be visualized in the medium (Figure 5C), often as aggregates around cellular debris. The capsid size and icosahedral morphology, as well as virion development within the cytoplasm is consistent with that of other isolated prasinoviruses [65]. This led us to conclude that the viral production of is the consequence of lysis of a subset of microalgae, defined as susceptible cells (S), and to dismiss the possibility of non-lytic virus production from budding or extrusion through the cell membrane of resistant microalga (R). Since the cultures are obtained from one single cell, this raised the question about the origin of the susceptible cells from a majority of virus-resistant microalga.

**Figure 5.**
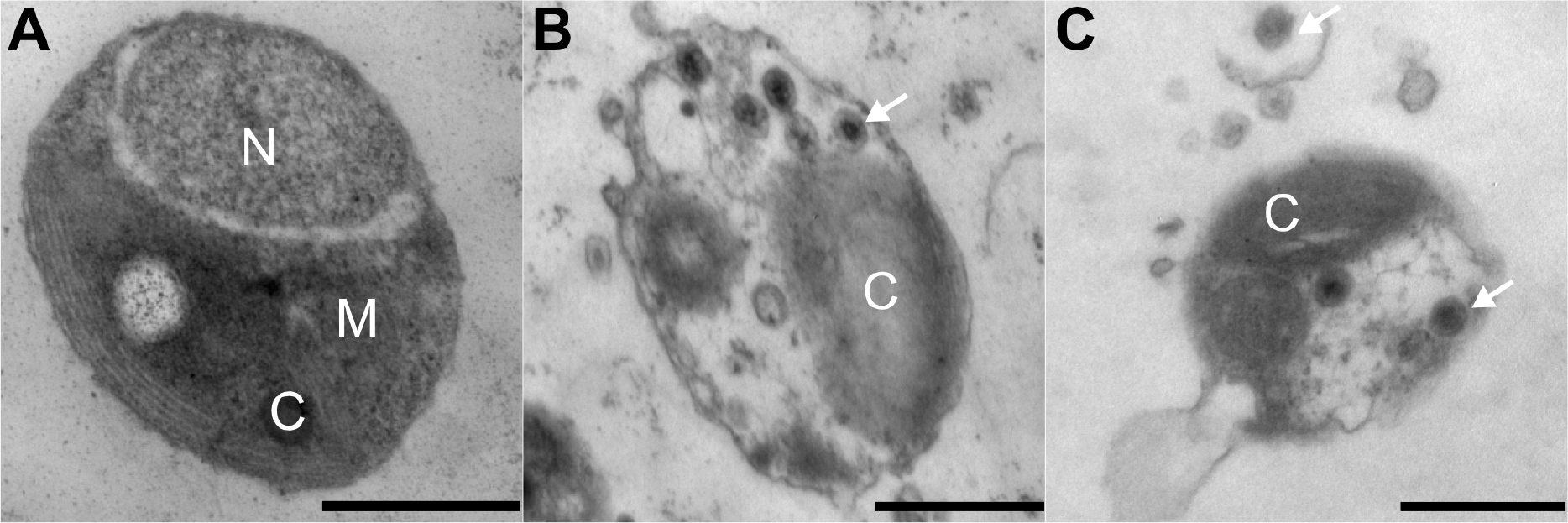
Transmission electron micrographs of *O. mediterraneus* RCC2590 cells from an actively growing culture. (A) The majority of cells have typical morphology while (B) 0.9% of cells contained visible viral particles, and (C) free viral particles were observed in the medium aggregated around infected cells. Scale bar-500 nm, C-chloroplast, M-mitochondrion, N-nucleus. White arrows indicate examples of virus particles.

### OmV2 susceptible (S) microalgae can be isolated from resistant cells (R), and *vice versa*

In order to test for the presence of both R and S phenotypes within cultures started from one single cell, we performed four independent experimental evolution studies by diluting cultures to single cells. Firstly, we produced a clonal virus free culture line by diluting the OmV2-producing RCC2590 culture to a single cell. This parent line (hereafter ML for “mother line”) was screened by PCR for OmV2 DNA and was found to be virus free. This demonstrated that virus-free cultures were easily obtainable by end-point dilution, presumably by random selection of an uninfected healthy individual cell. Secondly, distinct ML lines where used to generate hundreds of clones that were tested for susceptibility to OmV2 lysate. However, we cannot yet determine the phenotype at the level of a single cell, we can only test the phenotype of the culture, and thus the phenotype of the majority of the cells in a culture. The first ML line was sub-cloned via serial single cell bottlenecks every two weeks for 1 year in 24 mutation accumulation lines [66] and 3 out of 24 independent lines were visibly lysed upon addition of OmV2. Three independent experiments were performed over a shorter period where the ML was sub-cloned once by end-point dilution to generate 89, 46 and 69 clones from each of the parent ML lines, and tested for susceptibility to OmV2 lysate after 5 days. Visual inspection revealed no lysis in 159 clones while 45 clones lysed with addition of OmV2 (4, 15 and 26, from each experiment, respectively). This demonstrated that a minority of virus-free cultures, evolved from a single resistant cell, contained a majority of susceptible cells to OmV2 (24.9% ± 17.9%), and conversely, that the majority of clonal cultures contained a majority of cells resistant to lysis by OmV2.

To test whether cultures containing a majority of S cells similarly could generate an R phenotype, the three independent susceptible lines from the first mutation accumulation experimental evolution study were exposed to OmV2. Infected susceptible cultures showed a sharp decrease in cell concentration in the first 15 days, consistent with a lytic event dropping as low as 167 cells/ml, after which cultures recovered and cell concentrations reached that of uninfected controls (Supplementary Figure 4). These cultures subsequently showed no visible lysis when re-challenged with OmV2, demonstrating that S cultures can reliably switch to an R state.

### What are the genomic basis of the R to S switch?

The three OmV2 susceptible lines obtained from the experimental evolution assays were subsequently maintained in batch culture and intermittently tested over a year for susceptibility to OmV2, confirming that the S cultures contained a majority of susceptible cells. PFGE revealed that the SOC chromosome of the three OmV2-susceptible MA lines had experienced a visible size reduction, while no size change could be observed in the OmV2-resistant ML, which remained unchanged from the wild-type RCC2590 (Figure 6). One susceptible line, MA3, was selected for single molecule PacBio re-sequencing to elucidate the link to a switch from a virus-resistant to virus-susceptible state. This line was chosen as it had the smallest deletion in the SOC and could show the smallest change linked to a switch in phenotype. Resequencing showed that a 58 kb fragment was deleted from the end of the SOC (Figure 7). Surprisingly, the SOC comprised a mosaic of short repeated sequences mainly originating intrachromosomally, with some contributions from other chromosomes. Thus the deleted fragment was a pastiche of redundant SOC sequences, with only 18.7% of unique coding sequence deleted (6,153 bp) (Figure 7). The unique deleted genes included a nucleotide diphophosugar transferase, a transmembrane protein and a transposase, although the majority of genes had unknown functions. This functional profile was similar to that observed in the SOCs of other Mamiellales, which are enriched in hypothetical proteins, sugar modification and transporter genes, as well as mobile elements [13].

**Figure 6.**
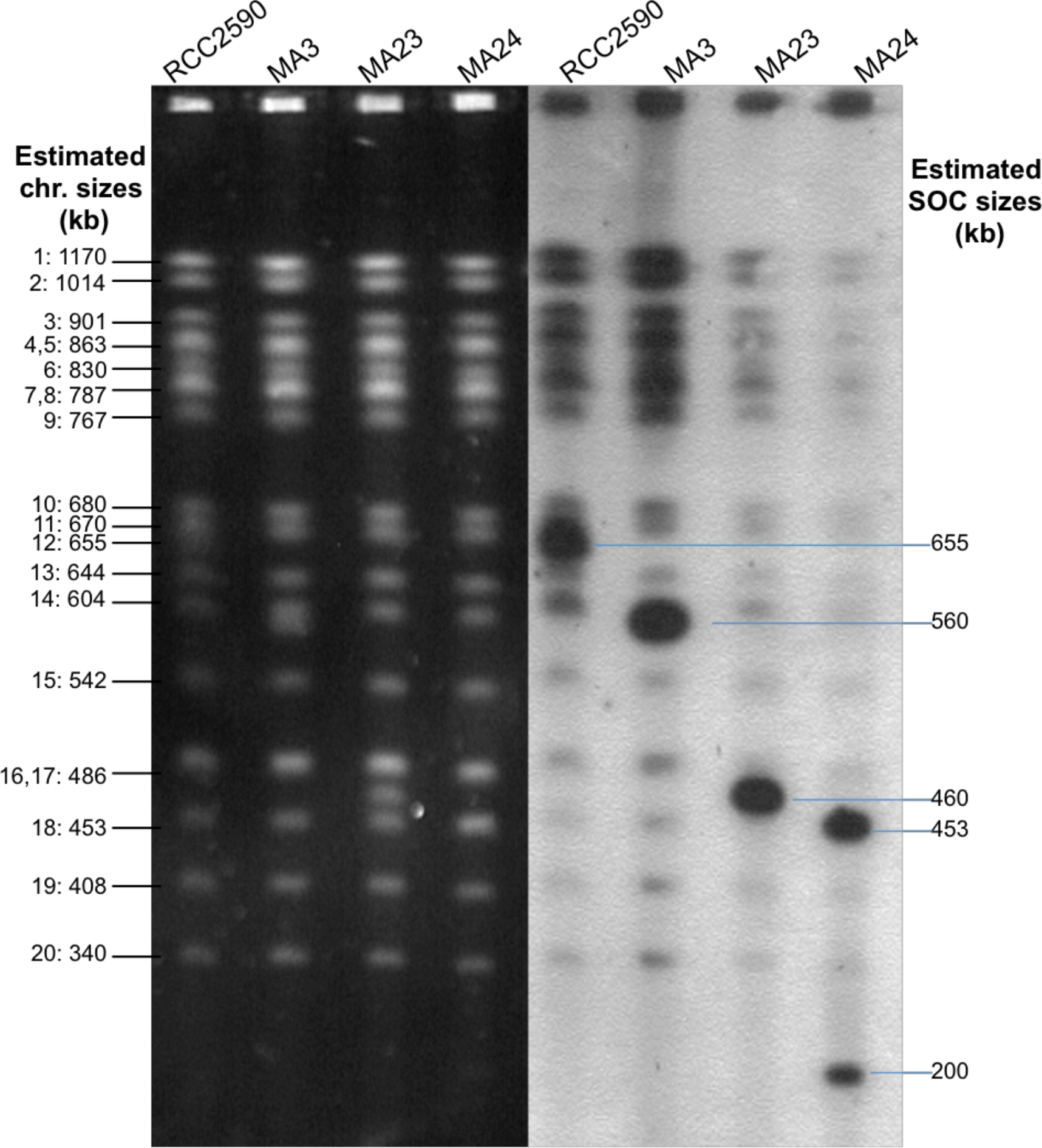
PFGE and hybridisation of *O. mediterraneus* OmV2-resistant parent RCC2590 and OmV2-susceptible (MA) lines. Left (black background): PFGE separation of chromosomes. Right (white background): in-gel hybridisation with SOC probes. Probes were generated by PCR using primer pairs listed in Supplementary Table 3, corresponding to SY2, SY3 and SY6.

**Figure 7.**
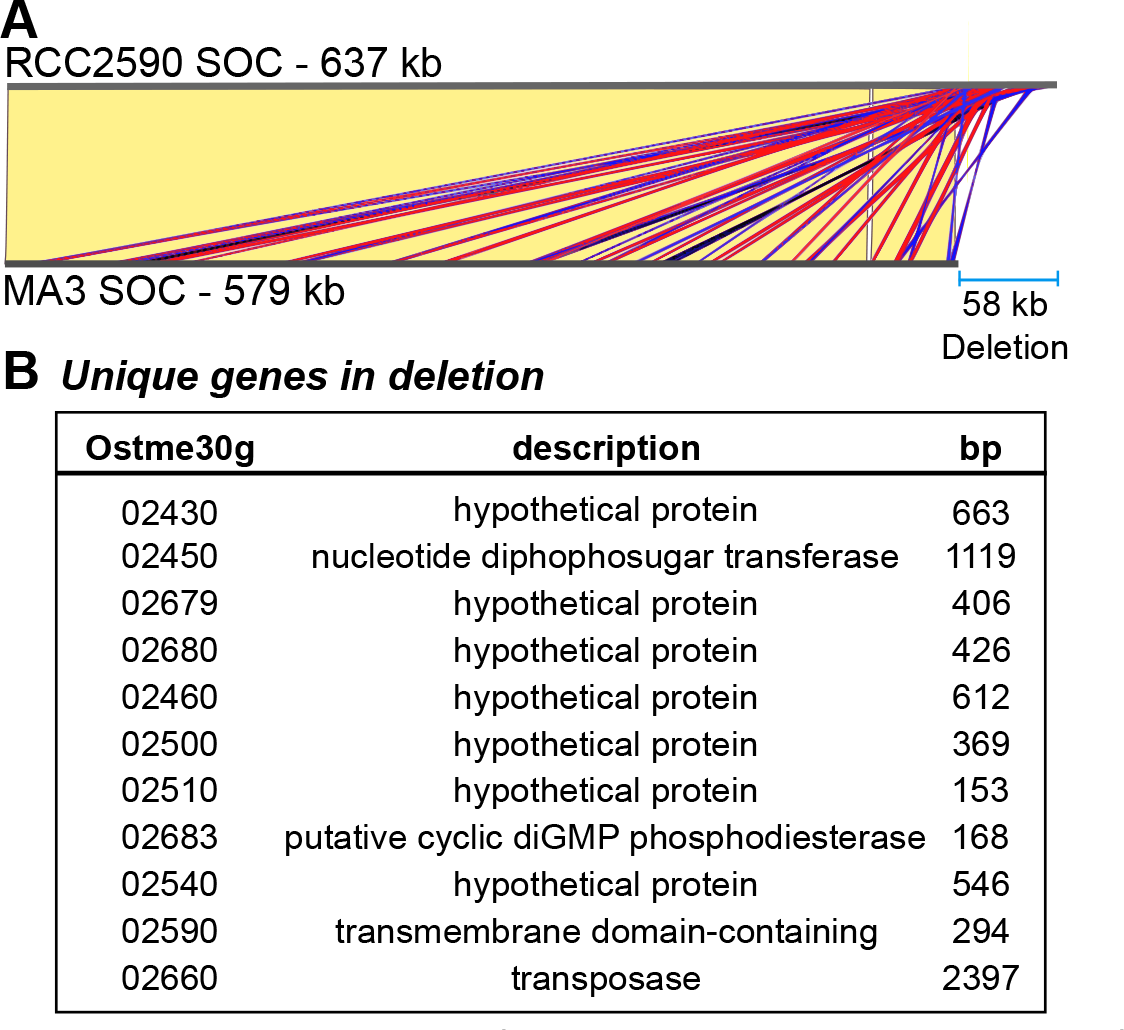
Comparison of Small Outlier Chromosome from *O. mediterraneus* RCC2590 and virus-susceptible MA3 line. (**A**) The MA3 SOC is virtually identical to the RCC2590 SOC (identical regions linked with yellow blocks) except for a 58 kb deletion at the end. The deletion consists of short sequences with significant identity to rest of the SOC, shown by red and blue blocks denoting sense and antisense matches, respectively. (**B**) Unique genes contained in the 58 kb region that were deleted in the MA3 SOC.

### Mathematical modelling of microalgae–virus dynamics

Our observations that a fraction of microalgae from a clonal culture was visibly infected, and that cultures derived from single R or S cells contained both phenotypes, were consistent with the existence of a phase switch within a clonal culture. The alternative hypothesis to a programmed phase switch, that is the generation of S cells from R cells by random spontaneous mutations, was not compatible with two observations. The first was the rapid emergence of resistance to prasinoviruses in the Mamiellophyceae. Clonal infected cultures typically restarted growth within 3 to 5 days post-infection, and reached pre-infection population sizes after 6 to 11 days post-infection [20][67]. Let us estimate the number of spontaneous mutations segregating in the culture at the time the virus is introduced. If *t*_*R*_ is the time required for recovery to pre-infection population density, in number of days, and provided post-infection growth rate can be approximated by pre-infection growth rate, the mutation leading to the resistant population appeared *t*_*R*_ days after starting the culture from one single cell. Given the typical growth rate of one division per day, the number of divisions in the culture when this mutation occurred can thus be approximated by 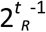. The rate of spontaneous mutations per genome, *U*_*G*_, in Mamiellophyceae has been estimated between 1×10^−2^ and 4×10^−3^ mutations per cell division [33]. The expected number of mutations at *t*_*R*_ in the microalga population is thus given by 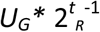, and the maximal estimate obtained with the largest *t*_*R*_ and *U*_*G*_ observed is 0.64. It is thus very likely that one (0.64) genome carrying one mutation was present in the culture at the time viruses were introduced. However, this spontaneous mutation had to confer viral resistance, which implies that 100% of spontaneous mutations confer resistance to viruses. The second observation was that resistance systematically occurred in all three S lines tested (Supplementary Figure 4). Furthermore, a previous study on the evolution of viral resistance in the sister species *O. tauri* and the prasinovirus OtV5 found all 38 independently infected cell lines became resistant to OtV5 within seven days [13]. This systematic resistance arousal again required 100% of spontaneous mutations to confer viral resistance, which is inconsistent with spontaneous mutations occurring randomly in the genome.

To investigate the consequences of such a phase switch on the microalga and virus population dynamics, we set up a host-virus interaction model of this system (Figure 8A). Susceptible hosts are infected by viral particles present in the environment, which ultimately lead to the lysis of the infected cell and the release of *v*_*S*_ new viral particles that survive in the environment at a rate *s*_*V*_. While reproducing asexually at rates *a*_*s*_ and *a*_*r*_, S algae produce a fraction *e*_*r*_ of R cells, while R algae generate a fraction *e*_*s*_ of S cells.

**Figure 8.**
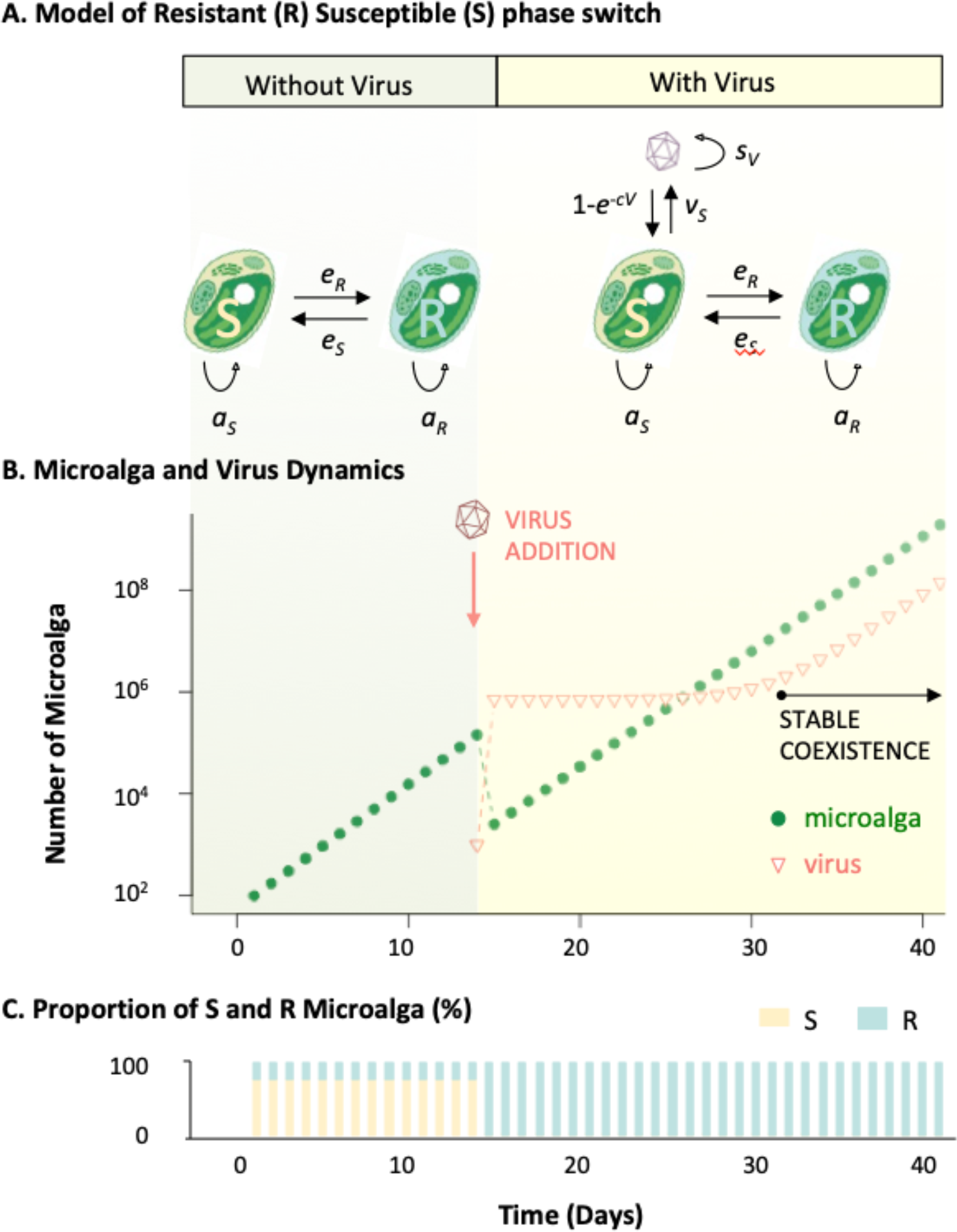
Model predictions of microalga-virus population dynamics as a function of model parameters. (A) Model parameters with and without virus. (B) Simulation of the evolution of the number of microalga cells during 40 days, viruses are introduced after 13 days and the system converged to its Virus/Microalga equilibrium behaviour after 18 days, *a*_*S*_ = 1.75, *a*_*R*_ = 1.7, *e*_*S*_ = *e*_*R*_ = 0.01. (C) Evolution of the proportion of Susceptible (S) and Resistant (R) microalga, initial S and R values have been set to reflect equilibrium proportion without virus.

In the absence of viruses, this model predicts the exponential growth typically observed in microalgae population dynamic experiments (Figure 8A#). The virus-free population growth rate converges toward an asymptotic value;

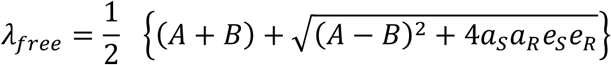

where *A*=(1 − *e*_*R*_)*a*_*S*_ and *B*=(1 − *e*_*S*_) *a*_*R*_ represent the rates at which S and R cells produce new born cells of the same kind as theirs. In such circumstances, the S/R ratio equals *e*_*S*_*a*_*R*_/(*λ*_*free*_ − *A*) which, for presumably small switching rates (*e*_*S*_, *e*_*R*_) and considering a cost to resistance, *i.e. a*_*S*_ > *a*_*R*_, leads to a larger fraction of S cells in the population, as shown in Figure 8B.

When the virus is introduced, the population size drops as S cells are eliminated from the population (Figure 8A), but only until the R cells, which were previously produced by the switch of S cells, take over the population (Figure 8B). During this transitory stage that only lasts for a few days, the S/R ratio changes and converges toward *e*_*S*_/(1 − *e*_*S*_), which for small values of *e*_*S*_, indicates that R cells dominate the population. The dynamics then enters a second asymptotic stage where the infection is maintained because R cells produce S cells that sustain the viral population (Figure 8A). The growth rate of such infected population quickly converges to;

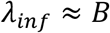

and the proportion of virus, *V*, to microalga, *M* (*M*=*S*+*R*), can then be approximated by:

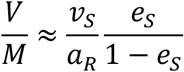

## Discussion

### Evolution of host and virus genomes

Analysis of the whole gene content of *O. mediterraneus* confirmed its phylogenetic position at the base of the *Ostreococcus* lineage. Maximum likelihood inference suggests that the *Ostreococcus* ancestor had a GC-poorer genome and contained a larger number of genes than the present species, revealing a global trend towards streamlining during the evolution of this lineage [69]. The comparison of the genome sequence of its resident prasinovirus, OmV2, with available prasinovirus sequence supports ancient divergence of OmV2 within the *Ostreococcus* infecting viruses for 13% of the genes, while the largest proportion of gene (36%) supports a recent divergence of OmV2 from *O. tauri* infecting viruses. Altogether, phylogenetic analysis of OmV2 genes dismisses co-evolution of this host–virus pair since the speciation of *O. mediterraneus*. Instead, it is consistent with substantial gene exchange between the ancestors of present day species-specific prasinoviruses [61].

Experimental evolution from single cells with and without virus enabled the existence of a phase switch between resistant (R) and susceptible (S) cells, and *vice versa*, to be revealed. Long read re-sequencing unveiled that S cells have a reduced Small Outlier Chromosome (SOC) size. Previous analyses of the *O. tauri* – OtV5 system have shown that *O. tauri* resistant lines tend to have larger SOC sizes [20] and increased expression of SOC-encoded genes associated primarily with carbohydrate modification and methylation [13]. We thus propose that the phase switch is associated to a hypermutation process involving a programmed rearrangement of the SOC chromosome as observed in S to R in *O. tauri* [13] and R to S changes (this study). The cell immunity toolkit is notorious for combining genetic and non-genetic mechanisms to produce phenotypic plasticity from a single genome [70]. Our knowledge about the antiviral cell immunity toolkit in microalga still looks very much like a blank page, and while we can only speculate about the genes involved in these rearrangements, our results support the proposed function of the SOC as a specialised chromosome involved in antiviral immunity in *Ostreococcus*. While all Mamiellales genomes sequenced to date possess a SOC [71][72][58][73], so called because it bears distinct genomic features — most notably a GC content approximately 5-15% lower than the rest of the genome, very few orthologous genes, with an estimated faster rate of evolution [74]— its role in antiviral resistance has been suggested in two species so far: *O. tauri* and *O. mediterraneus*. While there are no orthologous genes in the SOC for all sequenced Mamiellophyceae species to date, the same kinds of gene functions are overrepresented in the SOCs of all of these species, suggesting that variation in these functional genes imparts viral immunity in Mamiellophyceae. Genomic analyses of more distant relatives inside and outside the Archaeplastida are now needed to investigate whether this chromosome was present in an even more ancient eukaryotic ancestor.

### Population dynamics consequences

How does this phase switch translate into microalga–virus population dynamics? Mathematical modelling of the switch is qualitatively and quantitatively consistent with observed continuous virus production of OmV2 by *O. mediterraneus*. The continuous increase of viruses is the consequence of the production of novel viruses by susceptible cells, which are generated from resistant cells each generation. This model does not generate oscillatory dynamics after an initial lysis event, contrary to many previous host–lytic virus coexistence models [21]. For example, coexistence may result from an arms race between both partners, the underlying genetic mechanism being the continuously generation and fixation of novel mutations for resistance (host) and virulence (virus) [75], generating oscillatory dynamics [21]. Alternatively, oscillations may also be expected from frequency dependent mechanisms, where the frequency of infection is negatively associated to the virus prevalence, and may imply the regulation of gene expression of a virus virulence inhibitor [18].

The recently proposed ‘leaky resistance’ model from the bacteria–bacteriophage dynamic [76] does also predict continuous virus and host growth and could also be invoked to describe the R to S switch. Interestingly, the population collapse observed upon infection in culture is only predicted if susceptible microalga make up most of the population. This is predicted either if there is a bias between the S to R and the R to S switch (*e*_*S*_ > *e*_*R*_), or if susceptible microalgae have a higher growth rate than resistant microalgae, producing a trade-off between resistance and growth. While there is evidence of this trade-off in many bacterial–bacteriophage systems [77], there is little evidence for this in NCLDV resistant microalga yet. This might be due to the long generation time of microalga as compared to bacteria, but also because over longer time periods, our model predicts an equilibrium S/R ratio depending on four parameters, that are independent of the phenotype of the founder cell. Interestingly, long term competition experiments started with an equal number of S and R *O. tauri* cells resulted in the evolution of cultures with a majority of S cells [20], consistent with either a higher growth rate of S cells and thus a cost of resistance, or *e*_*S*_ > *e*_*R*_. In this system, the cost of resistance may be metabolic, as a consequence of an increase in the size of the outlier chromosome, which requires additional energy and resources for cell replication.

One fundamental property of a strategy is its evolutionary stability [78], which can be estimated by investigating whether a population composed of microalgae adopting such a strategy can be invaded by microalga playing alternative ones. Let us consider a microalga with a pure S strategy (it does not revert from S to R). This microalga cannot invade a culture with the resistant/susceptible switch strategy (hereafter RS strategy) in the presence of viruses, as the viruses will lyse it. It will however invade an RS culture in the absence of viruses, since microalga with a pure S strategy divide more rapidly than the R microalga. Now, if we consider a microalga with a pure R strategy, it will not invade a RS population in the absence of viruses, because of its lower growth rate than the S microalga. However, it will invade the RS population in the presence of viruses, as the R cells that are not reverted into S will invade the population. The RS strategy we describe is therefore not evolutionarily stable, neither in the presence nor the absence of viruses, so how did it persist in the long run? The most obvious explanation comes from the evolutionary theory of bet-hedging [79] that demonstrates that a strategy may be suboptimal under one environmental condition, but advantageous in the long run, when environmental conditions alternate randomly. The RS strategy outcompetes S microalga as soon as the environment contains a virus. The RS strategy will outcompete R microalga if virus presence and absence alternates [79]. A droplet of seawater can certainly experience changes in virus prevalence over short timescales, by mechanisms of marine virus decay [80][81], mixing of water masses or rain [82]. Last but not least, the RS strategy may confer a significant advantage to the microalga by lysing competitors. Indeed, strains propagating a virus have a higher probability of invasion in face of competitors that may be susceptible to the virus produced [83].

### What is the prevalence of the RS strategy in the environment?

The microalga–virus dynamics of the RS strategy predicts that the number of viruses increases linearly with the number of microalgae. However, it is difficult to predict how this single species dynamic translates at the community scale, when several microalga–virus pairs compete for the same resource [84]. A general prediction we may venture is that introducing RS microalgae in community models may increase the overall stability of the host–virus dynamics. This model is consistent with previously reported non ‘boom and bust’ dynamics in the natural environment [85][86], including persistent infection [87]. Within the simplified one microalga one virus experimental set up our model makes two predictions. First, it predicts the systematic arousal of resistance as a response to viral infection. This has been reported in many different species from diverged branches of the eukaryotic tree of life: Haptophyceae [18], Stramenopiles [88] and Chlorophytes [13]. Second, it predicts the steady production of viruses in apparently healthy cultures. While this has been explicitly reported in *Emiliania huxleyi*, *Phaeocystis pouchetii* [18] and *O. tauri* [20], it has also been observed in the dynamics of *Chlorella variabilis* [21]. Transcriptomes of several hundreds marine microalgae from the Marine Microbial Eukaryotic Transcriptome Sequencing Project (MMETSP) initiative revealed the expression of core NCDLV genes in 13 cultures [89]. While this may merely indicate infection, one of the haptophyte strains sequenced had been described as under persistent infection by a NCLDV [89]. Screening six *O. mediterraneus* transcriptomes in the MMETSP (Supplementary Table 2) for OmV2 sequences, we found evidence for a virus in strain RCC2596. This strain was isolated on the same day and site as the genome sequenced strain RCC2590 and, although closely related, they are non-identical possessing ~5 differences per 10,000 bp and different karyotypes [15]. RCC2596 was still available in culture in our lab and we could thus experimentally confirm the presence of viral DNA in the culture by PCR. Given that these strains were independently isolated, this implies stable coexistence was widespread in this population in the environment, but is simply not detected during the isolation of algae as the bulk culture grows reliably. We thus speculate that this phase switching is a common strategy within NCLDV-microalga interactions, in addition to mutation and selection mechanisms, that lead to pure resistant microalga strains and the arms-race type of interaction. This phase switch is the result of a yet to be described molecular mechanism of cell immunity in microalgae, mirroring the complexity of the NCLDV genomes they have been co-evolving with since the early stages of eukaryotic life [90].

## Accession numbers

The genome sequence for *Ostreococcus mediterraneus* RCC2590 and OmV2 can be found under Genbank accession numbers XX and YY. The complete gene annotations of RCC2590 can be accessed on ORCAE [34]. MMETSP sequence data can be accessed from https://www.imicrobe.us/#/projects/104. The genomes and their Genbank accession numbers used in this study are as follows: *Bathycoccus prasinos* (GCA_002220235.1), *Micromonas commoda* RCC299 (GCA_000090985.2), *Micromonas pusilla* CCMP1545 (GCA_000151265.1), *O. tauri* RCC4221 (CAID01000001.2 to CAID01000020.2, CR954200.2, CR954199.2), *O. lucimarinus* CCE9901 (GCA_000092065.1), BpV1 (HM004432.1), BpV2 (HM004430.1), MpV1 (HM004429.1), OtV1 (FN386611.1), OtV2 (FN600414.1), OtV5 (EU304328.2), OtV6 (JN225873.1), OlV1 (HM004431.1), OlV2 (KP874736.1), OlV3 (HQ633060.1), OlV4 (JF974316.1), OlV5 (HQ632827.1), OlV6 (HQ633059.1), OlV7 (KP874737.1), OmV1 (KP874735.1). *Ostreococcus* sp. RCC809 is available from the JGI genome portal under project ID 16233.

## Acknowledgements

We are grateful to the Genotoul bioinformatic platform for access to the computer cluster, the GeT sequencing platform for the PacBio sequencing, and the BIOPIC platform for access to the cytometers and microscopes. Our thanks to Alice Penrose and Yohan Gourbiere for their help in the manual annotation of *O. mediterraneus* and visual inspection of TEM screens.

## Author contribution

All molecular biology experiments on microalga and virus were performed at OOB (SY, MK, LB, SSB, ED, NG, HM). Illumina genome sequencing and assembly was performed at the Genoscope (JG, JMA). The genome annotation was completed at VIB (EV, SR) and corrected manually at OOB (MK, SY, NG, HM, GP, ED). MK and SY completed experimental evolution experiments. MLE performed TEM. Comparative genome analysis was performed by EV, KV, with help from FB and YD on OmV2 phylogeny, GC content evolution was performed by MG. SG completed mathematical model conception and analysis. SY, SG and GP wrote the manuscript, all authors contributed to manuscript editing.

## References

1. Short SM, Suttle CA. Sequence analysis of marine virus communities reveals that groups of related algal viruses are widely distributed in nature. Appl Environ Microbiol 2002; 68: 1290–1296.

2. Laber CP, Hunter JE, Carvalho F, Collins JR, Hunter EJ, Schieler BM, et al. Coccolithovirus facilitation of carbon export in the North Atlantic. Nat Microbiol 2018; 3: 537–547.

3. Short SM. The ecology of viruses that infect eukaryotic algae. Environ Microbiol 2012; 14: 2253–2271.

4. Hingamp P, Grimsley N, Acinas SG, Clerissi C, Subirana L, Poulain J, et al. Exploring nucleo-cytoplasmic large DNA viruses in Tara Oceans microbial metagenomes. ISME J 2013; 7: 1678–1695.

5. Wilson WH, Van Etten JL, Allen MJ. The Phycodnaviridae: The Story of How Tiny Giants Rule the World. Curr Top Microbiol Immunol 2009; 328: 1–42.

6. Mateus MD. Bridging the Gap between Knowing and Modeling Viruses in Marine Systems—An Upcoming Frontier. Front Mar Sci 2017; 3: 284.

7. Suttle CA. Marine viruses”-major players in the global ecosystem. Nat Rev Microbiol 2007; 5: 801–812.

8. Brussaard CPD, Wilhelm SW, Thingstad F, Weinbauer MG, Bratbak G, Heldal M, et al. Global-scale processes with a nanoscale drive: the role of marine viruses. ISME J 2008; 2: 575–578.

9. Tragin M, Vaulot D. Green microalgae in marine coastal waters: The Ocean Sampling Day (OSD) dataset. Sci Rep 2018; 8: 14020.

10. Weynberg KD, Allen MJ, Wilson WH. Marine Prasinoviruses and Their Tiny Plankton Hosts: A Review. Viruses 2017; 9.

11. Derelle E, Ferraz C, Escande ML, Eychenie S, Cooke R, Piganeau G, et al. Life-cycle and genome of OtV5, a large DNA virus of the pelagic marine unicellular green alga Ostreococcus tauri. PLoS One 2008; 3: e2250.

12. Derelle E, Monier A, Cooke R, Worden AZ, Grimsley NH, Moreau H. Diversity of Viruses Infecting the Green Microalga Ostreococcus lucimarinus. J Virol 2015; 89: 5812–5821.

13. Yau S, Hemon C, Derelle E, Moreau H, Piganeau G, Grimsley N. A Viral Immunity Chromosome in the Marine Picoeukaryote, Ostreococcus tauri. PLoS Pathog 2016; 12: e1005965.

14. Blanc-Mathieu R, Krasovec M, Hebrard M, Yau S, Desgranges E, Martin J, et al. Population genomics of picophytoplankton unveils novel chromosome hypervariability. Sci Adv 2017; 3: e1700239.

15. Subirana L, Péquin B, Michely S, Escande M-L, Meilland J, Derelle E, et al. Morphology, Genome Plasticity, and Phylogeny in the Genus Ostreococcus Reveal a Cryptic Species, O. mediterraneus sp. nov. (Mamiellales, Mamiellophyceae). Protist 2013; 164: 643–659.

16. Erez Z, Steinberger-Levy I, Shamir M, Doron S, Stokar-Avihail A, Peleg Y, et al. Communication between viruses guides lysis-lysogeny decisions. Nature 2017; 541: 488–493.

17. Dunigan DD, Fitzgerald LA, Van Etten JL. Phycodnaviruses: a peek at genetic diversity. Virus Res 2006; 117: 119–132.

18. Thyrhaug R, Larsen A, Thingstad F, Bratbak G. Stable coexistence in marine algal host-virus systems. Mar Ecol Prog Ser 2003; 254: 27–35.

19. Van Etten JL, Lane LC, Meints RH. Viruses and viruslike particles of eukaryotic algae. Microbiol Rev 1991; 55: 586–620.

20. Thomas R, Grimsley N, Escande M, Subirana L, Derelle E, Moreau H. Acquisition and maintenance of resistance to viruses in eukaryotic phytoplankton populations: Viral resistance in Mamiellales. Environ Microbiol 2011; 13: 1412–1420.

21. Frickel J, Sieber M, Becks L. Eco-evolutionary dynamics in a coevolving host-virus system. Ecol Lett 2016; 19: 450–459.

22. Derelle E, Ferraz C, Lagoda P, Eychenie S, Cooke R, Regad F, et al. Dna libraries for sequencing the genome of Ostreococcus tauri (Chlorophyta, Prasinophyceae): The smallest free-living eukaryotic cell. J Phycol 2002; 38: 1150–1156.

23. Wohl T, Brecht M, Lottspeich F, Ammer H. The use of genomic DNA probes for in-gel hybridization. Electrophoresis 1995; 16: 739–741.

24. Winnepenninckx B, Backeljau T, De Wachter R. Extraction of high molecular weight DNA from molluscs. Trends Genet TIG 1993; 9: 407.

25. Gnerre S, Maccallum I, Przybylski D, Ribeiro FJ, Burton JN, Walker BJ, et al. High-quality draft assemblies of mammalian genomes from massively parallel sequence data. Proc Natl Acad Sci U S A 2011; 108: 1513–1518.

26. Robbens S, Khadaroo B, Camasses A, Derelle E, Ferraz C, Inze D, et al. Genome-wide analysis of core cell cycle genes in the unicellular green alga Ostreococcus tauri (vol 22, pg 589, 2005). Mol Biol Evol 2005; 22: 1158–1158.

27. Kearse M, Moir R, Wilson A, Stones-Havas S, Cheung M, Sturrock S, et al. Geneious Basic: an integrated and extendable desktop software platform for the organization and analysis of sequence data. Bioinforma Oxf Engl 2012; 28: 1647–1649.

28. Chin C-S, Alexander DH, Marks P, Klammer AA, Drake J, Heiner C, et al. Nonhybrid, finished microbial genome assemblies from long-read SMRT sequencing data. Nat Methods 2013; 10: 563–569.

29. Blanc-Mathieu R, Verhelst B, Derelle E, Rombauts S, Bouget F-Y, Carré I, et al. An improved genome of the model marine alga Ostreococcus tauri unfolds by assessing Illumina de novo assemblies. BMC Genomics 2014; 15: 1103.

30. Foissac S, Bardou P, Moisan A, Cros M-J, Schiex T. EUGENE’HOM: A generic similarity-based gene finder using multiple homologous sequences. Nucleic Acids Res 2003; 31: 3742–3745.

31. Keeling PJ, Burki F, Wilcox HM, Allam B, Allen EE, Amaral-Zettler LA, et al. The Marine Microbial Eukaryote Transcriptome Sequencing Project (MMETSP): illuminating the functional diversity of eukaryotic life in the oceans through transcriptome sequencing. PLoS Biol 2014; 12: e1001889.

32. Gremme G, Kurtz S, Brendel V, Sparks. Engineering a software tool for gene structure prediction in higher organisms. Inf Softw Technol 2005; 47: 965–978.

33. Kim D, Langmead B, Salzberg SL. HISAT: a fast spliced aligner with low memory requirements. Nat Methods 2015; 12: 357.

34. Sterck L, Billiau K, Abeel T, Rouzé P, Van de Peer Y. ORCAE: online resource for community annotation of eukaryotes. Nat Methods 2012; 9: 1041.

35. Kearse M, Moir R, Wilson A, Stones-Havas S, Cheung M, Sturrock S, et al. Geneious Basic: an integrated and extendable desktop software platform for the organization and analysis of sequence data. Bioinforma Oxf Engl 2012; 28: 1647–1649.

36. Lowe TM, Eddy SR. tRNAscan-SE: a program for improved detection of transfer RNA genes in genomic sequence. Nucleic Acids Res 1997; 25: 955–964.

37. Vandepoele K, Van Bel M, Richard G, Van Landeghem S, Verhelst B, Moreau H, et al. pico-PLAZA, a genome database of microbial photosynthetic eukaryotes. Environ Microbiol 2013; 15: 2147–2153.

38. Altschul SF, Gish W, Miller W, Myers EW, Lipman DJ. Basic local alignment search tool. J Mol Biol 1990; 215: 403–10.

39. Enright AJ, Van Dongen S, Ouzounis CA. An efficient algorithm for large-scale detection of protein families. Nucleic Acids Res 2002; 30: 1575–1584.

40. Li L, Stoeckert CJ, Roos DS. OrthoMCL: identification of ortholog groups for eukaryotic genomes. Genome Res 2003; 13: 2178–2189.

41. Edgar RC. MUSCLE: multiple sequence alignment with high accuracy and high throughput. Nucleic Acids Res 2004; 32: 1792–1797.

42. Stamatakis A. RAxML version 8: a tool for phylogenetic analysis and post-analysis of large phylogenies. Bioinforma Oxf Engl 2014; 30: 1312–1313.

43. Felsenstein J. PHYLIP (Phylogeny Inference Package) version 3.6. 2005.

44. Katoh K, Kuma K, Toh H, Miyata T. MAFFT version 5: improvement in accuracy of multiple sequence alignment. Nucleic Acids Res 2005; 33: 511–518.

45. Ronquist F, Teslenko M, van der Mark P, Ayres DL, Darling A, Hohna S, et al. MrBayes 3.2: efficient Bayesian phylogenetic inference and model choice across a large model space. Syst Biol 2012; 61: 539–542.

46. Ranwez V, Harispe S, Delsuc F, Douzery EJP. MACSE: Multiple Alignment of Coding SEquences accounting for frameshifts and stop codons. PloS One 2011; 6: e22594.

47. Criscuolo A, Gribaldo S. BMGE (Block Mapping and Gathering with Entropy): a new software for selection of phylogenetic informative regions from multiple sequence alignments. BMC Evol Biol 2010; 10: 210.

48. Price MN, Dehal PS, Arkin AP. FastTree 2--approximately maximum-likelihood trees for large alignments. PloS One 2010; 5: e9490.

49. Yang Z, Nielsen R. Mutation-selection models of codon substitution and their use to estimate selective strengths on codon usage. Mol Biol Evol 2008; 25: 568–579.

50. Hershberg R, Petrov DA. Evidence That Mutation Is Universally Biased towards AT in Bacteria. PLoS Genet 2010; 6: e1001115.

51. Romiguier J, Ranwez V, Douzery EJP, Galtier N. Contrasting GC-content dynamics across 33 mammalian genomes: Relationship with life-history traits and chromosome sizes. Genome Res 2010; 20: 1001–1009.

52. Lassalle F, Périan S, Bataillon T, Nesme X, Duret L, Daubin V. GC-Content Evolution in Bacterial Genomes: The Biased Gene Conversion Hypothesis Expands. PLoS Genet 2015; 11: e1004941.

53. Dutheil J, Boussau B. Non-homogeneous models of sequence evolution in the Bio++ suite of libraries and programs. BMC Evol Biol 2008; 8: 255.

54. Gueguen L, Gaillard S, Boussau B, Gouy M, Groussin M, Rochette NC, et al. Bio++: efficient extensible libraries and tools for computational molecular evolution. Mol Biol Evol 2013; 30: 1745–1750.

55. Yang Z, Kumar S, Nei M. A new method of inference of ancestral nucleotide and amino acid sequences. Genetics 1995; 141: 1641–1650.

56. Marie D, Rigaut-Jalabert F, Vaulot D. An improved protocol for flow cytometry analysis of phytoplankton cultures and natural samples. Cytom Part J Int Soc Anal Cytol 2014; 85: 962–968.

57. Krasovec M, Eyre-Walker A, Grimsley N, Salmeron C, Pecqueur D, Piganeau G, et al. Fitness Effects of Spontaneous Mutations in Picoeukaryotic Marine Green Algae. G3 GenesGenomesGenetics 2016; 6: 2063–2071.

58. Worden AZ, Lee J-H, Mock T, Rouzé P, Simmons MP, Aerts AL, et al. Green evolution and dynamic adaptations revealed by genomes of the marine picoeukaryotes Micromonas. Science 2009; 324: 268–272.

59. Robbens S, Derelle E, Ferraz C, Wuyts J, Moreau H, Van de Peer Y. The complete chloroplast and mitochondrial DNA sequence of Ostreococcus tauri: organelle genomes of the smallest eukaryote are examples of compaction. Mol Biol Evol 2007; 24: 956–968.

60. Weynberg KD, Allen MJ, Ashelford K, Scanlan DJ, Wilson WH. From small hosts come big viruses: the complete genome of a second Ostreococcus tauri virus, OtV-1. Environ Microbiol 2009; 11: 2821–2839.

61. Moreau H, Piganeau G, Desdevises Y, Cooke R, Derelle E, Grimsley N. Marine prasinovirus genomes show low evolutionary divergence and acquisition of protein metabolism genes by horizontal gene transfer. J Virol 2010; 84: 12555–63.

62. Bellec L, Clerissi C, Edern R, Foulon E, Simon N, Grimsley N, et al. Cophylogenetic interactions between marine viruses and eukaryotic picophytoplankton. BMC Evol Biol 2014; 14: 59.

63. Clerissi C, Grimsley N, Desdevises Y. Genetic exchanges of inteins between prasinoviruses (phycodnaviridae). Evol Int J Org Evol 2013; 67: 18–33.

64. Monier A, Claverie JM, Ogata H. Horizontal gene transfer and nucleotide compositional anomaly in large DNA viruses. BMC Genomics 2007; 8: 456.

65. Derelle E, Ferraz C, Escande M-L, Eychenié S, Cooke R, Piganeau G, et al. Life-cycle and genome of OtV5, a large DNA virus of the pelagic marine unicellular green alga Ostreococcus tauri. PloS One 2008; 3: e2250.

66. Krasovec M, Eyre-Walker A, Grimsley N, Salmeron C, Pecqueur D, Piganeau G, et al. Fitness Effects of Spontaneous Mutations in Picoeukaryotic Marine Green Algae. G3 Bethesda Md 2016; 6: 2063–2071.

67. Yau S, Hemon C, Derelle E, Moreau H, Piganeau G, Grimsley N. A Viral Immunity Chromosome in the Marine Picoeukaryote, Ostreococcus tauri. PLoS Pathog 2016; 12: e1005965.

68. Krasovec M, Eyre-Walker A, Sanchez-Ferandin S, Piganeau G. Spontaneous mutation rate in the smallest photosynthetic eukaryotes. Mol Biol Evol 2017.

69. Wolf YI, Koonin EV. Genome reduction as the dominant mode of evolution. Bioessays 2013; 35: 829–837.

70. Meyers LA, Bull JJ. Fighting change with change: adaptive variation in an uncertain world. Trends Ecol Evol; 17: 551–557.

71. Derelle E, Ferraz C, Rombauts S, Rouzé P, Worden AZ, Robbens S, et al. Genome analysis of the smallest free-living eukaryote Ostreococcus tauri unveils many unique features. Proc Natl Acad Sci U S A 2006; 103: 11647–11652.

72. Palenik B, Grimwood J, Aerts A, Rouze P, Salamov A, Putnam N, et al. The tiny eukaryote Ostreococcus provides genomic insights into the paradox of plankton speciation. Proc Natl Acad Sci U A 2007; 104: 7705–10.

73. Moreau H, Verhelst B, Couloux A, Derelle E, Rombauts S, Grimsley N, et al. Gene functionalities and genome structure in Bathycoccus prasinos reflect cellular specializations at the base of the green lineage. Genome Biol 2012; 13: R74.

74. Jancek S, Gourbière S, Moreau H, Piganeau G. Clues about the genetic basis of adaptation emerge from comparing the proteomes of two Ostreococcus ecotypes (Chlorophyta, Prasinophyceae). Mol Biol Evol 2008; 25: 2293–2300.

75. Gandon S, Buckling A, Decaestecker E, Day T. Host-parasite coevolution and patterns of adaptation across time and space. J Evol Biol 2008; 21: 1861–1866.

76. Chaudhry WN, Pleska M, Shah NN, Weiss H, McCall IC, Meyer JR, et al. Leaky resistance and the conditions for the existence of lytic bacteriophage. PLoS Biol 2018; 16: e2005971.

77. Lennon JT, Khatana SAM, Marston MF, Martiny JBH. Is there a cost of virus resistance in marine cyanobacteria? ISME J 2007; 1: 300–312.

78. Maynard Smith J. Evolution and the Theory of Games. 1982. Cambridge University Press.

79. Philippi T, Seger J. Hedging One’s Evolutionary Bets, Revisited. Trends Ecol Evol 1989; 4: 41–44.

80. Suttle CA, Chen F. Mechanisms and rates of decay of marine viruses in seawater. Applied and Environmental Microbiology 1992; 58: 3721–3729.

81. Mojica KDA, Brussaard CPD. Factors affecting virus dynamics and microbial host-virus interactions in marine environments. FEMS Microbiol Ecol 2014; 89: 495–515.

82. Sharoni S, Trainic M, Schatz D, Lehahn Y, Flores MJ, Bidle KD, et al. Infection of phytoplankton by aerosolized marine viruses. Proc Natl Acad Sci 2015; 112: 6643.

83. Brown SP, Le Chat L, De Paepe M, Taddei F. Ecology of microbial invasions: amplification allows virus carriers to invade more rapidly when rare. Curr Biol CB 2006; 16: 2048–2052.

84. Winter C, Bouvier T, Weinbauer MG, Thingstad TF. Trade-Offs between Competition and Defense Specialists among Unicellular Planktonic Organisms: the “Killing the Winner” Hypothesis Revisited. Microbiol Mol Biol Rev MMBR 2010; 74: 42–57.

85. Coolen MJL. 7000 years of Emiliania huxleyi viruses in the Black Sea. Science 2011; 333: 451–452.

86. Moniruzzaman M, Wurch LL, Alexander H, Dyhrman ST, Gobler CJ, Wilhelm SW. Virus-host relationships of marine single-celled eukaryotes resolved from metatranscriptomics. Nat Commun 2017; 8: 16054.

87. Short SM, Short CM. Quantitative PCR reveals transient and persistent algal viruses in Lake Ontario, Canada. Environ Microbiol 2009; 11: 2639–2648.

88. Tarutani K, Nagasaki K, Yamaguchi M. Viral impacts on total abundance and clonal composition of the harmful bloom-forming phytoplankton Heterosigma akashiwo. Appl Environ Microbiol 2000; 66: 4916–4920.

89. Gallot-Lavallée L, Blanc G. A Glimpse of Nucleo-Cytoplasmic Large DNA Virus Biodiversity through the Eukaryotic Genomics Window. Viruses 2017; 9: 17.

90. Koonin EV, Senkevich TG, Dolja VV. The ancient Virus World and evolution of cells. Biol Direct 2006; 1: 29.

