## Supplementary Material for "Virus-host coexistence in phytoplankton through the genomic lens"

#### Supplemental Material

**Supplementary Figure 1.** Colinearity between *O. mediterraneus* RCC2590 (x-axis) and *O. tauri* RCC4221 (y-axis) chromosomes.

**Supplementary Figure 2.** Phylogenomic analysis of *O. mediterraneus* within the Archaeplastida based on 188 single gene families.

**Supplementary Figure 3.** Reconstructed ancestral GC contents from first codon (left), second codon (middle) and third codon (right) positions.

**Supplementary Figure 4.** Growth curves of uninfected control and virus infected *O. mediterraneus* culture lines.

**Supplementary Table 1.** OmV2 predicted protein coding genes with available functional annotation.

**Supplementary Table 2.** *O. mediterraneus* strains and genomic and transcriptomic datasets used in this study.

**Supplementary Table 3.** Primers used in this study on *O. mediterraneus* RCC2590 and OmV2.

**Supplementary Table 4***.* Definition of the parameters of the model and numerical estimations from litterature.

**Supplementary Table 5***.* Species list and genome versions used for annotation and comparative genomics analysis.

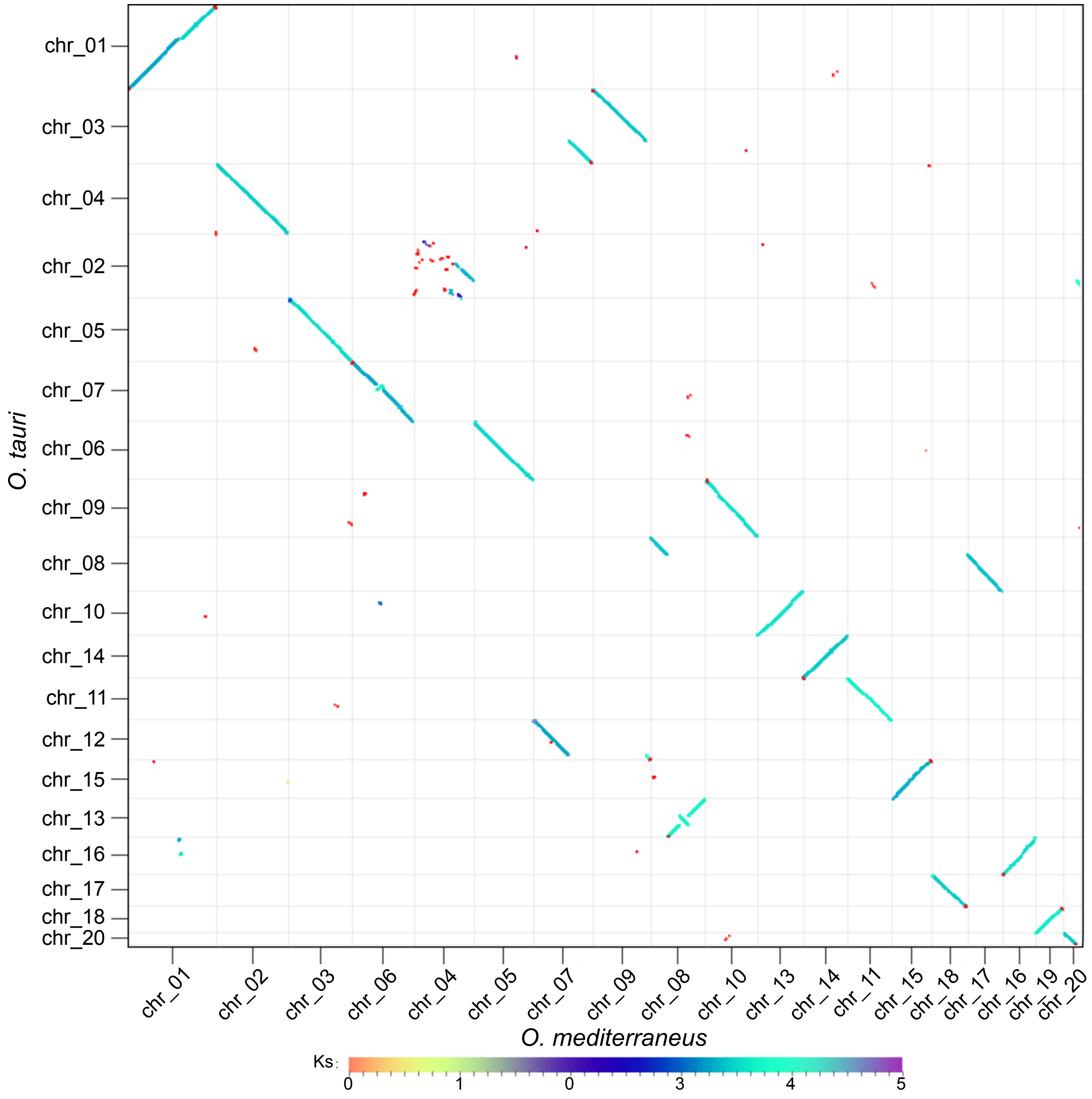


**Supplementary Figure 1.** Colinearity between *O. mediterraneus* RCC2590 (x-axis) and *O. tauri* RCC4221 (y-axis) chromosomes. Note: The small outlier chromosomes (chromosome 19 of *O. tauri* and chromosome 12 of *O. mediterraneus*) do not share colinearity and are not shown.

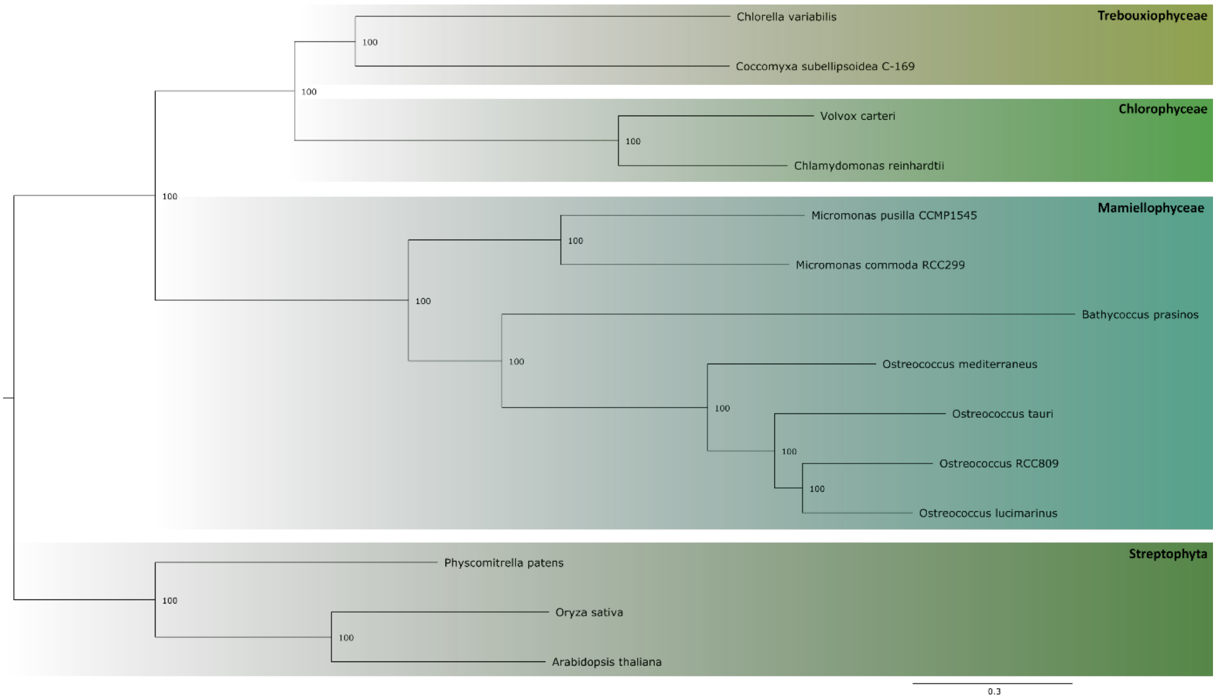


**Supplementary Figure 2.** Phylogenomic analysis of *O. mediterraneus*. Tree generated using RaxML (model PROTGAMMAWAG, 100 bootstraps) from the concatenated amino acid sequences of 188 single copy gene families shared in the Chlorophyta.

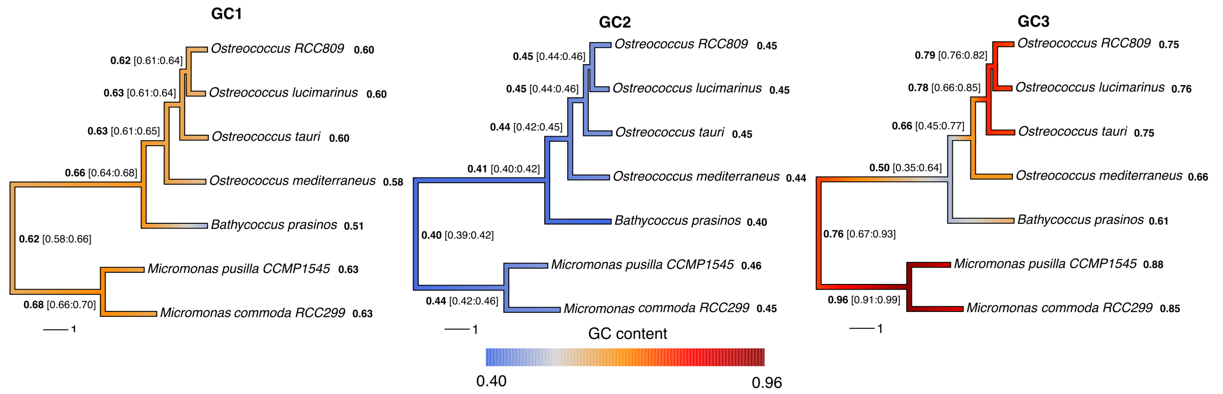


**Supplementary Figure 3.** Reconstructed ancestral GC contents from first codon (left), second codon (middle) and third codon (right) positions are shown on the same scale with position specific GC contents of extant genomes shown after the species names in bold and the ancestral estimates shown at the nodes with ranges in square brackets.

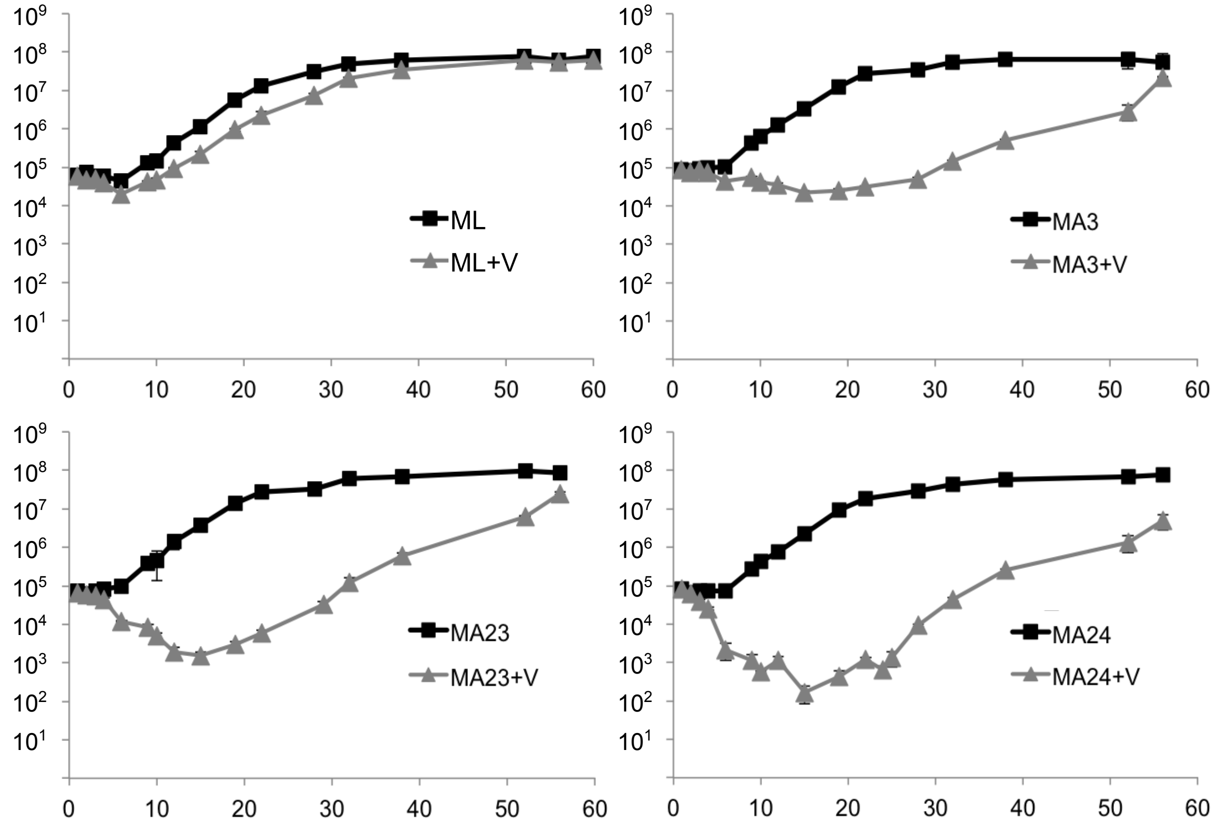


**Supplementary Figure 4.** Growth curves of uninfected control and virus infected *O. mediterraneus* culture lines. Cell concentrations ($ml^{-1}$) are showed in the ordinate and time (days) on the abscissa. Black squares – uninfected controls, Grey triangles – OmV2 infected. Error bars show the standard deviation between three replicate culture flasks. ML is a virus-free OmV2 resistant line, MA3, MA23 and MA24 are OmV2-susceptible lines.

### Supplementary Tables

**Supplementary Table 1.** OmV2 predicted protein coding genes with available functional annotation. BBH : Best Blast Hit.

| CDS | Gene names and genomic features | BBH | CDS | Gene names and genomic features | BBH |
|---|---|---|---|---|---|
| | **DNA, RNA, replication, recombination and repair** | | | **Lipid/Fatty acid metabolism** | |
| **orf00008** | Exonuclease | OmV1 | **orf00172** | Patatin-like phospholipase | OtV1 |
| **orf00020** | ATPase | OmV1 | | **Protein synthesis, modification and degradation** | |
| **orf00075** | Poxvirus A22 protein | OlV4 | **orf00009** | Zinc knuckle | OmV1 |
| **orf00077** | Thymidylate synthase complementing protein | OmV1 | **orf00018** | Peptidase family M41 | OlV7 |
| **orf00151** | Proliferating cell nuclear antigen, N-terminal domain | OtV5 | **orf00052** | DegT/DnrJ/EryC1/StrS aminotransferase family | OlV2 |
| **orf00163** | NTPase/helicase | OtV2 | **orf00087** | NAD(P)-binding Rossmann-like domain | OmV1 |
| **orf00166** | NTPase/helicase | OtV2 | **orf00128** | WLM domain | OlV2 |
| **orf00183** | YqaJ-like viral recombinase domain | OlV1 | **orf00252** | Clp protease | OtV2 |
| **orf00228** | D5 N terminal like | OtV1 | **orf00266** | Proline dehydrogenase | OlV1 |
| **orf00235** | RNase H | OtV5 | | **Signaling** | |
| **orf00243** | Eukaryotic DNA topoisomerase I | OmV1 | **orf00032** | Ring finger domain | OlV5 |
| **orf00251** | ATP dependent DNA ligase domain | OtV2 | **orf00209** | Protein kinase domain | OtV5 |
| **orf00253** | RsgA GTPase | OtV5 | **orf00295** | Rhodanese-like domain | OlV3 |
| **orf00304** | DNA polymerase family B | OtV5 | | **Structural** | |
| | **DNA methylation and site-specific endonucleases** | | **orf00091** | Large eukaryotic DNA virus major capsid protein | OmV1 |
| **orf00090** | Type III restriction enzyme, res subunit | OlV1 | **orf00101** | Major capsid protein N-terminus | OtV1 |
| **orf00131** | D12 class N6 adenine-specific DNA methyltransferase | OtV1 | **orf00114** | Large eukaryotic DNA virus major capsid protein | OtV2 |
| **orf00160** | C-5 cytosine-specific DNA methylase | OlV4 | **orf00133** | Major capsid protein N-terminus | OlV2 |
| **orf00180** | SWIB/MDM2 domain | OtV5 | **orf00233** | Large eukaryotic DNA virus major capsid protein | OlV1 |
| | **Sugar manipulation enzymes** | | **orf00302** | Large eukaryotic DNA virus major capsid protein | OtV5 |
| **orf00015** | GDP-mannose 4,6 dehydratase | OtV1 | **orf00307** | Large eukaryotic DNA virus major capsid protein | OlV2 |
| **orf00048** | Glycosyl transferase family 2 | OtV1 | | **Miscellaneous** | |
| **orf00053** | GDP-mannose 4,6 dehydratase | OlV3 | **orf00010** | T5orf172 domain | OlV1 |
| **orf00241** | Phosphofructokinase | OlV4 | **orf00021** | cell division protein | OlV4 |
| **orf00246** | Glycosyltransferase family 25 (LPS biosynthesis protein) | OtV1 | **orf00023** | Methyltransferase domain | OlV1 |
| | **Transcription** | | **orf00044** | PhoH-like protein | OtV1 |
| **orf00036** | Transcription factor S-II (TFIIS) | OtV1 | **orf00049** | Thiamine pyrophosphate enzyme, N-terminal TPP binding domain | OtV2 |
| **orf00126** | mRNA capping enzyme | OlV3 | **orf00071** | Baculoviridae p74 conserved region | No hit |
| **orf00186** | SNF2 family N-terminal domain | OtV5 | **orf00073** | Chaperone of endosialidase | No hit |
| **orf00154** | Poxvirus Late Transcription Factor VLTF3 like | OlV2 | **orf00079** | ABC1 family | OlV1 |
| **orf00201** | Transcription factor TFIID (or TATA-binding protein, TBP) | OtV1 | **orf00111** | Chaperone of endosialidase | OlV7 |
| **orf00217** | Transcription factor TFIIB repeat | OtV1 | **orf00118** | NUDIX domain | OtV5 |
| **orf00220** | mRNA capping enzyme, catalytic domain | OmV1 | **orf00159** | Cytidylyltransferase-like | OtV2 |
| **orf00221** | mRNA capping enzyme, beta chain | OtV2 | **orf00162** | 2OG-Fe(II) oxygenase superfamily | OtV1 |
| | **Nucleotide metabolism** | | **orf00171** | viralA-type inclusion protein | OlV3 |
| **orf00182** | Ribonuclease-III-like | OtV2 | **orf00195** | MazG nucleotide pyrophosphohydrolase domain | OlV4 |
| **orf00191** | Ribonucleotide reductase, barrel domain | OmV1 | **orf00237** | 2OG-Fe(II) oxygenase superfamily | OmV1 |
| **orf00215** | Ribonucleotide reductase, small chain | OmV1 | **orf00240** | Phytanoyl-CoA dioxygenase (PhyH) | OtV6 |
| **orf00250** | 5' nucleotidase, deoxy (Pyrimidine), cytosolic type C protein (NT5C) | OlV4 | **orf00244** | A nuclease family of the HNH/ENDO VII superfamily with conserved AHH | OtV5 |
| **orf00275** | Thymidine kinase | OlV7 | **orf00248** | PQ loop repeat | OtV6 |
| **orf00284** | dUTPase | OlV1 | **orf00287** | 2OG-Fe(II) oxygenase superfamily | OlV1 |
| | | | **orf00289** | Domain of unknown function (DUF814) | OtV6 |
| | | | **orf00291** | 2OG-Fe(II) oxygenase superfamily | OtV2 |

**Supplementary Table 2.** *O. mediterraneus* strains and genomic and transcriptomic datasets used in this study. RCC2590 is genomic sequence data and strains with an MMETSP identifier are transcriptome sequences. Abbreviations: N/A, non-applicable; PE, paired-end; MP, mate-paired; RCC, Roscoff Culture Collection; MA3, MA23 and MA24 (Mutation Accumulation lines derived from RCC2590); MMETSP, Marine Microbial Eukaryotic Transcriptome Sequencing Project; assem., nuclear genome assembly; aln reads, read fragments re-aligned to genome; ID, identifier. *percentage of properly aligned PE reads to the reference RCC2590 genome.

| Strain ID | MMETSP ID | isolation date | site | coordinates | reads | assem. (Mb) | aln reads* (%) |
|---|---|---|---|---|---|---|---|
| RCC2590 | N/A | 23 Mar. 2009 | Thau | 43°24'N 3°36'E | 10,257,955 PE 13,542,963 MP 80× PACBIO | 13.86 | 85 |
| MA3 | N/A | N/A | N/A | N/A | 747,558 PE 80× PACBIO | 13.81 | 75 |
| MA16 | N/A | N/A | N/A | N/A | N/A | N/A | N/A |
| MA23 | N/A | N/A | N/A | N/A | N/A | N/A | N/A |
| MA24 | N/A | N/A | N/A | N/A | N/A | N/A | N/A |
| RCC2572 | 0929 | 16 Jun. 2006 | Bages | 43°03'14"N 2°59'54"E | 43,616,308 PE | 11.6 | 83 |
| RCC1621 | 0930 | 2 Jun. 2003 | SOLA | 42°29'18"N 3°08'42"E | 29,821,828 PE | 12.4 | 89 |
| RCC2596 | 0932 | 23 Mar. 2009 | Thau | 43°24'N 3°36'E | 27,255,407 PE | 14.0 | 95 |
| RCC2573 | 0936 | 18 Jan. 2008 | Leucate | 42°48'24"N 3°01'27"E | 28,940,482 PE | 12.2 | 89 |
| RCC2593 | 0937 | 23 Mar. 2009 | Thau | 43°24'N 3°36'E | 25,147,103 PE | 11.5 | 84 |
| RCC1107 | 0938 | 1 Jan. 2006 | Bages | 43°03'14"N 2°59'54"E | 27,277,118 PE | 12.9 | 88 |

**Supplementary Table 3.** Primers used in this study on *O. mediterraneus* RCC2590 and OmV2. Abbreviations: Seq. F, forward primer sequence; Seq. R, reverse primer sequence; SOC, small outlier chromosome; BOC, big outlier chromosome; DNApol, viral DNA polymerase B; MCP, viral Major Capsid Protein.

| Primer ID | Seq. F. (5'–3') | Seq. R. (5'–3') | Size (bp) | Target | Reference |
|---|---|---|---|---|---|
| SY1F/R | TCGAACACGAGGACTTAGCG | ATCAGCAGGGTTGTCATCCG | 448 | SOC | This study |
| SY2F/R | CACTACGTCACCGCCGATAA | CTTCACACTTTGTGCCCGTG | 517 | SOC | This study |
| SY3F/R | CATCATGTGCGCCTTTCTCG | GAAACAAGCACAACGTCCCC | 642 | SOC | This study |
| SY4F/R | ATGGATTCCGAGGAACCGTG | GAACGCACCCGATCCAGTTA | 640 | OmV2 | This study |
| SY5F/R | CCTCCCACGACGTTTCTTCA | CGCACTGTAATACGCACACG | 603 | BOC | This study |
| SY6F/R | GCGTGAACGCATCGACAATC | CCGGATACCCAAACCGTTGA | 534 | SOC | This study |
| SY7F/R | ATCAATGCGATTCGTTGCGG | ATTGCCGAGAGTGATGCCAA | 654 | OmV2 | This study |
| 18SF/R | ACCTGGTTGATCCTGCCAG | TGATCCTTCCGCAGGTTCAC | 1765 | 18S rRNA | Grimsley *et al.*, 2010 |
| VpolAS4/ VpolAAS1 | GARGGIGCIACIGTIYTNGA | CCIGTRAAICCRTAIACISWRTTCAT | 320 | DNApol | Clerissi *et al.*, 2014 |
| VmcpAS3/ VmcpAAS1 | GGIGGICARMGIRTIGAYAA | TGI ACY TGY TCD ATI ARR TAY TCR TG | 350 | MCP | Clerissi *et al.*, 2014 |

**Supplementary Table 4**. Definition of the parameters of the model and numerical estimations from litterature. *nd* : no data. The numerical parameter values used to draw Figure 8 are indicated in bold.

| | Parameter Definition | Value | Host/Virus system | Reference |
|---|---|---|---|---|
| $v_S$ | Burst size : number of viruses produced by each lysed cell | 5-35<br>**5** | *O. tauri* RCC4221 / OtV5<br>O. *mediterraneus* / OmV2 | [20]; this study<br>**Fig. 8** |
| $e_S$ | Proportion of resistant cells reverting into susceptible state | 0.007-0.07<br>**0.01** | *O. mediterraneus* RCC2590/ OmV2 | This study<br>**Fig. 8** |
| $e_R$ | Proportion of susceptible cells switching to resistant state | 0.05<br><br>**0.01** | *O. tauri* RCC4221/ OtV5 | Inferred from data in [17]<br>**Fig. 8** |
| $s_V$ | Survival of the viral particle in the environment | 1<br>**1** | OmV2 | This study<br>**Fig. 8** |
| $a_S$ | Growth rate of Susceptible cells (without the virus) | 1.9<br>1.95 ± 0.14<br>**1.75** | MA3 line | [78]; this study<br>**Fig. 8** |
| $a_R$ | Growth rate of resistant cells | 1.75<br>1.92 ± 0.17<br>**1.7** | RCC2590 without virus<br>average of 21 lines | [78]; this study<br>**Fig. 8** |
| $c$ | Microalga virus encounter rate | Nd<br>**0.9** | | **Fig. 8** |

**Supplementary Table 5**. Species list and genome versions used for annotation and comparative genomics analysis. For species denoted with asterisk functional annotations (GO annotations and InterPro domains) were retrieved using the Uniprot Gene Association File (downloaded 10/09/2015). For all other species InterPro was ran (January 2016) and mapped to GO terms.

| **Species** | **Source** | **PubmedID** |
|---|---|---|
| *Aureococcus anophagefferens* | JGI 1.0 | 21368207 |
| *Asterochloris sp. Cgr/DA1pho v2.0* | JGI 7.45.13 | / |
| *Auxenochlorella protothecoides* | Beijing Genomics Institute 1.0 | 25012212 |
| *Arabidopsis thaliana ** | TAIR10 | 11130711 |
| *Amborella trichopoda ** | Amborella v1 | 24357323 |
| *Bathycoccus prasinos* | Ghent University | 22925495 |
| *Chondrus crispus* | ENSEMBL protists release 28 | 23536846 |
| *Caenorhabditis elegans ** | ENSEMBL release 81 | 9851916 |
| *Cyanidioschyzon merolae* | Tokyo University | 15071595 |
| *Chlorella variabilis NC64A* | JGI 1.0 | 20852019 |
| *Chlamydomonas reinhardtii* | JGI 5.5 (Phytozome 10.2) | 17932292 |
| *Coccomyxa subellipsoidea C-169* | JGI 2.0 (Phytozome 10.2) | 22630137 |
| *Dictyostelium discoideum ** | ENSEMBL protist release 28 | 15875012 |
| *Drosophila melanogaster** | ENSEMBL release 81 | 10731132 |
| *Emiliania huxleyi ** | ENSEMBL protist release 28 | 23760476 |
| *Ectocarpus siliculosus* | Ghent University | 20520714 |
| *Fragilariopsis cylindrus* | JGI 1.0 | 28092920 |
| *Galdieria sulphuraria* | ENSEMBL protists release 28 | 23471408 |
| *Homo sapiens ** | ENSEMBL release 81 | 11181995 |
| *Helicosporidium sp.* | Illinois University 1.0 | 24809511 |
| *Micromonas pusilla strain CCMP1545* | Ghent University | 24273312 |
| *Micromonas sp RCC299* | JGI 3.0 | 19359590 |
| *Mus musculus ** | ENSEMBL release 81 | 12466850 |
| *Nannochloropsis gaditana ** | ENSEMBL protist release 28 | 23966634 |
| *Ostreococcus lucimarinus* | JGI 2.0 | 17460045 |
| *Ostreococcus sp RCC809* | JGI 2.0 | / |
| *Oryza sativa ** | MSU RGAP 7 | 16100779 |
| *Ostreococcus tauri* | Ghent University v2.0 | 25494611 |
| *Phytophtora sojae** | ENSEMBL protist release 28 | 16946065 |
| *Physcomitrella patens* | Phytozome 9.1 (v1.6) | 18079367 |
| *Picochlorum costavermella* | This study | / |
| *Picochlorum sp. SENEW3 (SE3)* | Rutgers University 1.0 | 24965277 |
| *Paramecium tetraurelia ** | ENSEMBL protist release 28 | 17086204 |
| *Phaeodactylum tricornutum ** | ASM15095v2 | 18923393 |
| *Saccharomyces cerevisiae strain S288C ** | ENSEMBL release 81 | 8849441 |
| *Schizosaccharomyces pombe ** | ENSEMBL fungi release 28 | 11859360 |
| *Thalassiosira pseudonana* | JGI 3.0 | 15459382 |
| *Volvox carteri* | JGI 2.0 (Phtyozome 10.2) | 20616280 |
